# Pangenome of U.S. ex-PVP and Wild Sorghum Reveals Structural Variants and Selective Sweeps Shaping Adaptation and Trait Improvement

**DOI:** 10.1101/2025.07.25.666886

**Authors:** Justine K. Kitony, Emily Murray, Kelly Colt, Ryan Lynch, Nick Allsing, Nolan T. Hartwick, Tiffany Duong, Jocelyn Saxton, Nadia Shakoor, Todd P. Michael

## Abstract

Sorghum is a cereal crop grown for food, feed, and biofuel, yet the genomic diversity of elite commercial lines remains underexplored. Here, we assembled a sorghum pangenome from long-read sequences of 46 U.S. ex-Plant Variety Protection Act (ex-PVP) cultivars and a haplotype-resolved assembly of one wild accession. The pangenome revealed largely conserved genomic architecture across elite lines but identified presence–absence variations (PAVs) in 28% of gene families, including genes involved in stress responses and starch metabolism, such as *STARCH BRANCHING ENZYME I (SBE1)*. Selective sweep analyses detected strong signals at key flowering-time loci (*Ma1/SbPRR37, Ma2, Ma6*) and circadian/light regulators (*PHOT1, DET1, XAP5*), with *PHYA* showing copy number variations (CNVs). Analysis of 24 wild accessions, beyond the core pangenome panel, detected CNVs in *MULTIDRUG AND TOXIC COMPOUND EXTRUSION (MATE)* transporters, potentially reflecting selection or breeding for enhanced herbicide detoxification. These results highlight breeding-shaped regions and position the pangenome as a resource for sorghum improvement.

## Main

Sorghum (*Sorghum bicolor*) is a C_4_ monocot renowned for its exceptional resilience to drought, heat, and low-fertility soils ^1^. As the fifth most important cereal crop globally, it is cultivated for diverse purposes—including food, animal feed, alcoholic beverages, and biofuel—driven by its nutritional value (high protein, essential micronutrients, gluten-free) and adaptability to abiotic stresses. These attributes make sorghum a vital crop for enhancing food security and promoting sustainable agriculture in the face of climate change ^1–5^.

Sorghum, originating in Africa, was introduced to the U.S. in the late 19^th^ century through germplasm collections of sweet, grain, and sudangrass types ^6–9^. Early public breeding programs played a key role in improvement, with notable successes such as the development of the drought-tolerant ‘Atlas’ variety in 1936 ^8^. The discovery of cytoplasmic male sterility (CMS) and corresponding fertility restorer (Rf) genes in the mid-20^th^ century enabled commercial hybrid production, boosting yields by up to 40% by the 1950s ^7,10^.

From the 1960s to 1980s, U.S. sorghum breeding shifted toward the private sector. Companies like Pioneer Hi-Bred began developing proprietary hybrids protected for 20 years under the 1970 Plant Variety Protection Act (PVPA). These hybrids featured dwarf stature (suitable for mechanical harvesting), early maturity, stay-green traits (e.g., SC35), and resistance to anthracnose, greenbug, downy mildew, and smut ^7,11^. As a result, the United States emerged as the world’s leading exporter of sorghum, accounting for approximately 15% of global production ^12–14^

Most genomic studies in sorghum have relied on the ‘BTx623’ reference genome ^1,15,16^, which represents a single grain-type line and fails to capture the full extent of genetic variation within cultivated germplasm. This reference bias limits the discovery of presence–absence variations (PAVs), SVs, and CNVs that underlie important agronomic traits ^17–19^. Pangenome approaches provide a broader framework for characterizing SVs across diverse accessions ^20,21^. Short-read-based pangenomes in sorghum have identified 35,719 genes, with ~53% showing PAVs and 79 loci absent from the reference genome ^22,23^. Long-read-based pangenomes have uncovered even greater diversity—44,079 gene families across 16 accessions, including 222.6 Mb of novel sequence ^21^. Similarly, high-quality assemblies from ten bioenergy sorghum genotypes uncovered thousands of SVs, including those associated with traits like iron transport ^24^. Accessibility of long-read sequencing (e.g., PacBio HiFi, ONT) is now enabling detection of novel gene content and SVs associated with traits ^25,26^.

From early domestication to modern crop improvement, selection—both natural and artificial— has left footprints across plant genomes. These genomic signatures reveal how crops have been shaped for key agronomic traits across time and environments. In wheat, genomic scans have uncovered loci controlling flowering time ^27^; in rice, genes conferring pathogen resistance ^28^; and in maize and cotton, variants linked to yield and stress tolerance ^29,30^. Sorghum is no exception. Selective sweeps have been reported at loci such as *SHATTERING1 (SH1)*, which affects seed dispersal, and the *DRY NAC* transcription factor, involved in sugar accumulation ^31–33^. Beyond selection on classical domestication and yield-related genes, natural and artificial selection has also acted on sorghum’s internal time-of-day (TOD) regulatory network (circadian rhythms), which governs key developmental processes such as vegetative growth and reproductive timing ^34–36^. These rhythms also influence the optimal timing of agricultural inputs and support sorghum’s adaptation across broad latitudinal ranges. Selection at the *Ma1* locus, encoding *PSEUDO-RESPONSE REGULATOR 37 (SbPRR37)*, exemplifies adaptation to temperate climates and remains a target in modern breeding for bioenergy ^37^. Yet many regions under selection remain unresolved due to the underrepresentation of elite commercial germplasm genomes and the complexity of sorghum’s breeding history ^38–40^.

Here we present a long-read-based sorghum pangenome constructed from 46 U.S. ex-PVP lines and wild accessions, revealing approximately 28% PAVs in gene families, localized SVs, selection signatures in TOD loci, and CNVs associated with key agronomic traits. This resource enhances our understanding of sorghum adaptation and provides a foundation for genomic characterization and crop improvement through modern breeding approaches ^39,41,42^.

## Results

### High-Quality Genome Assemblies of Ex-PVP and Wild Sorghum Accessions

We assembled high-quality genomes for 46 elite ex-PVP lines and both haplotypes of a wild sorghum accession, ‘PI156549.’ Genome sizes ranged from 610 to 779 megabases (Mb), with confirmed gene models ranging from 30,000 to 33,000 per genome (Fig. 1a). The new assemblies achieved levels of contiguity comparable to the ‘BTx623’ reference ^16^, with N50 values reaching up to 19 Mb (Fig. 1b, Supplementary Fig. 1a-c, Supplementary Table 1), reflecting reference-quality assemblies across the dataset.

**Fig. 1.**
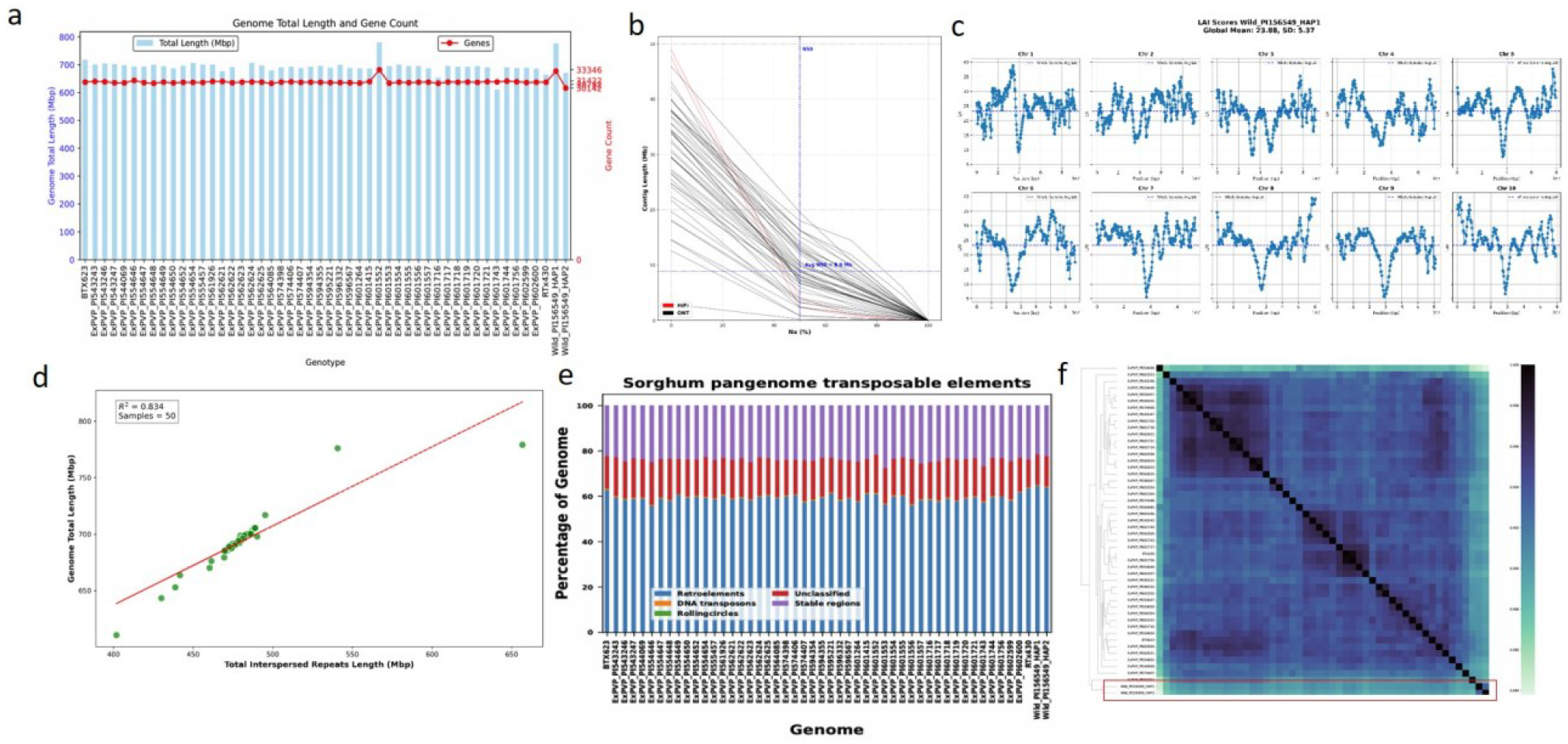
Genome Assembly Metrics, Transposable Elements (TEs), and Genome Architecture of the U.S. Ex-PVP and Wild Sorghum Pangenome. **a**, Genome length (Mb; right y-axis) and gene count (left y-axis) across assembled accessions. **b**, Assembly contiguity measured by N50 values (Mb). **c**, Example LTR Assembly Index (LAI) distribution across chromosomes for the wild accession ‘PI156549’ Haplotype 1, with a global mean of 23.88 and a standard deviation (SD) of 5.37. **d**, Correlation between total interspersed repeat content and genome size. **e**, TEs classes expressed as a percentage of genome size. Colors represent TEs categories as shown in the legend. **f**, Clustered heatmap of adjacency values based on pairwise K-mer comparisons between genomes. The ex-PVP lines exhibit largely conserved nucleotide identity, forming a homogeneous cluster distinct from the wild accession haplotypes (red box). Pangenome accessions include elite ex-PVP lines and wild sorghum samples. Full sample and sequencing details are in Supplementary Table 1; source data are available in the associated data file.

Assembly quality was assessed using the LTR Assembly Index (LAI), with scores ranging from 20 to 30, consistent with reference-grade assemblies (Fig. 1c, Supplementary Table 1). BUSCO analysis confirmed high completeness of both genome assemblies (92.60–99.10% complete BUSCOs) and predicted proteins (91.70–98.50%), supporting an accurate gene space view across diverse genotypes (Supplementary Table 2).

Total interspersed repeat content showed a strong correlation with genome size (R^2^ = 0.834; Fig. 1d), underscoring the dominant role of repeats in genome expansion. Centromeric tandem repeat expansions also correlated with genome size (R^2^ = 0.747; Supplementary Fig. 1d). Transposable elements (TEs), which make up ~65% of each genome assembly (Fig. 1e), dominate sorghum genome composition and have broadly shaped its structure, regulation, and adaptability ^43^. Among these, retroelements (LTRs) were the most abundant class, followed by DNA transposons and rolling-circle elements. Notably, a substantial proportion (~15–20%) of the TEs content remained unclassified across genomes, even when using high-resolution annotations from Helixer ^44^. This unclassified fraction highlights the presence of highly divergent or lineage-specific elements that escape conventional TEs classification approaches ^45,46^.

Analysis based on K-mer profiles revealed genetic clustering among the ex-PVP lines, with some accessions showing closer relatedness but overall forming a cohesive group. The wild accession formed a distinct outgroup (Fig. 1f), consistent with its contribution to novel K-mers in the PanKmer collector’s curve, which shows an open pangenome (Supplementary Fig. 1e, Supplementary Table 3).

### Sorghum Pangenome Captures Gene Diversity Missed by the BTx623 Reference

Using the constructed pangenome, we characterized gene content variation that would be overlooked when relying solely on the ‘BTx623’ reference^16^. In total, 36,004 gene families (orthogroups) were identified (Supplementary Table 4), comprising 71.5% core genes, 2.2% soft-core genes, 16.7% dispensable genes, and 9.6% private genes (Fig. 2a). Gene accumulation curves showed continued pangenome growth and core genome decline, approaching a plateau that suggests most gene diversity was captured (Fig. 2b). A power law fit of novel gene family discovery (a = 4.13, R^2^ = 0.999) confirmed a highly closed sorghum pangenome (Fig.2c) ^47^. Comparison to the ‘BTx623’ reference revealed 6,389 additional gene families, many absent from the reference but present in wild and elite ex-PVP lines (Fig. 2d). Gene PAVs analysis across the 50 genomes revealed shared gene family patterns among the ex-PVPs (Fig. 2e), with genespace visualization further highlighting overall gene conservation across chromosomal segments (Fig. 2f.

The 6,389 pangenome exclusive orthogroups (Fig. 2d) include key KEGG functions absent from the reference ‘BTx623’ (Fig. 2g), such as chloroplast electron transport components—ATP SYNTHASE SUBUNIT BETA (ATPB), NADH DEHYDROGENASE SUBUNIT H (NDHH), and NADH DEHYDROGENASE SUBUNIT F (NDHF); stress response regulators—GLUTATHIONE S-TRANSFERASE TAU 1 (GSTU1) and ABSCISIC ALDEHYDE OXIDASE 3 (AAO3); and specialized metabolism genes—FLAVONOID 3’-HYDROXYLASE (F3’H), SALICYLIC ACID CARBOXYL METHYLTRANSFERASE (SAMT) and TREHALOSE-6-PHOSPHATE SYNTHASE 1 (TPS1). Notably detected is STARCH BRANCHING ENZYME I (SBE1), which controls starch branching and may harbor allelic variants preferentially retained in ex-PVP lines for improved grain quality or digestibility traits ^48^. In total, 982 KEGG orthology (KOs) associations were identified across these orthogroups (Supplementary Table 5). These KOs suggest hidden variation in energy metabolism, detoxification, and hormone-mediated stress adaptation, echoing findings in other crops where pangenome analyses revealed adaptive traits beyond reference genomes ^49^. These findings also demonstrate the value of the pangenome in uncovering agronomically relevant diversity that is missed by single-reference approaches ^50^.

**Fig. 2.**
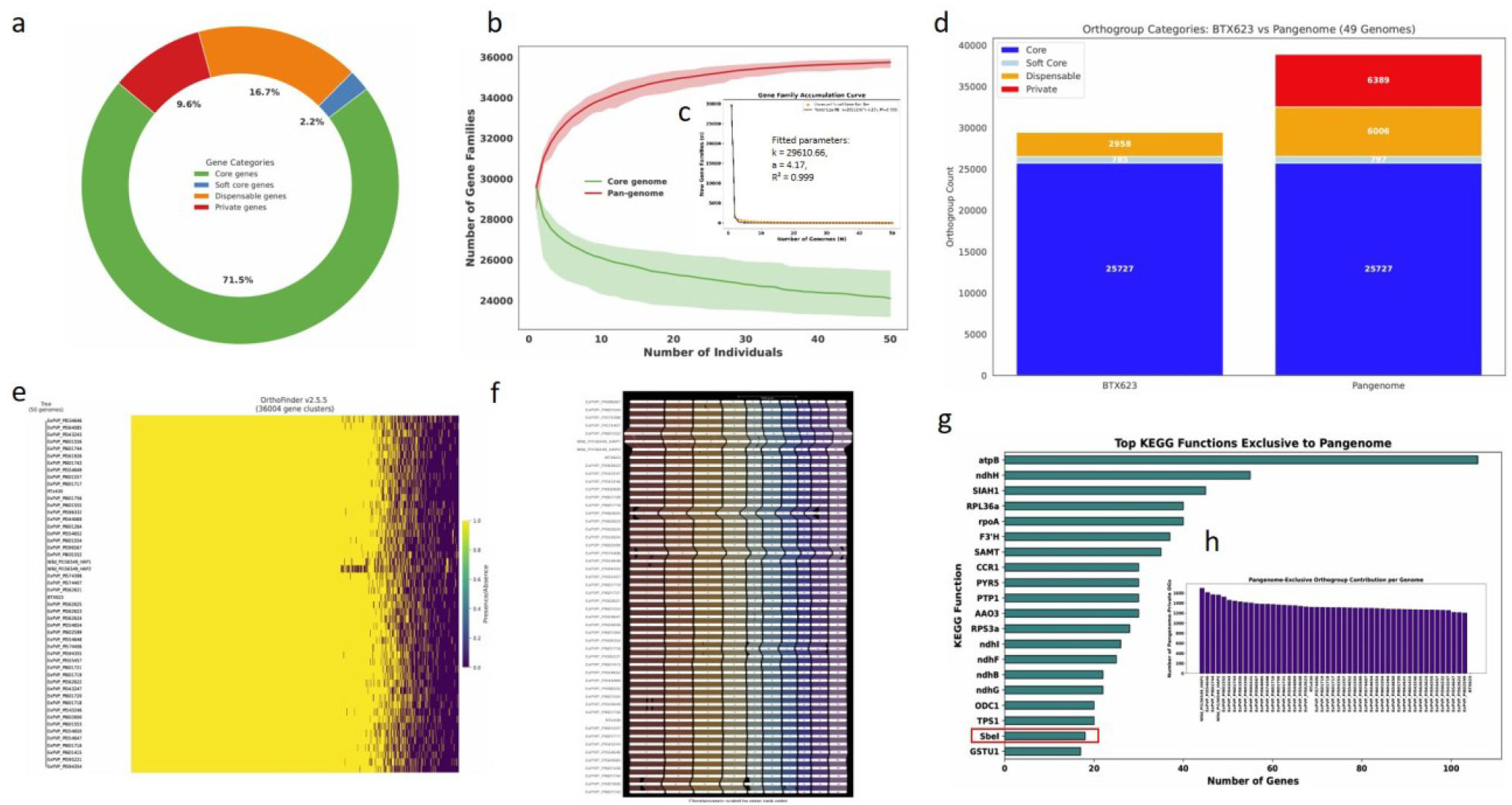
Pangenome-Wide Gene Presence–Absence Variation Reveals Genes Missing from the BTx623 Reference. **a**, Classification of 36,004 gene families into core (green), soft-core (blue), dispensable (orange), and private (red). **b**, Collectors curves showing pan- and core-genome orthogroups dynamics across genomes sampled. **c**, Power law fit of novel gene family discovery indicating a closed sorghum pangenome as evidenced by the plateauing collectors curve **d**, Stacked bar plot comparing gene categories in the ‘BTx623’ reference (left) and the full pangenome (right), the red bar reveals 6,389 additional gene families present in the pangenome. **e**, Gene family presence–absence variation across accessions highlights shared patterns among ex-PVP lines. Yellow indicates presence; black indicates absence. **f**, Genespace riparian plot illustrating macro-synteny across sorghum genomes, revealing broad genic similarity and a mostly conserved structure with a few notable rearrangements. **g**, a bar plot of enriched KEGG orthology (for genes in pangenome-exclusive orthogroups). The orthologies include ATP synthase subunit beta (atpB), NADH dehydrogenase subunits (ndhH, ndhI, ndhF, ndhB, ndhG), E3 ubiquitin-protein ligase SIAH1 (SIAH1), ribosomal protein L36a (RPL36a), DNA-directed RNA polymerase subunit alpha (rpoA), Flavonoid 3’-hydroxylase (F3’H), Salicylic acid carboxyl methyltransferase (SAMT), Cinnamoyl-CoA reductase 1 (CCR1), Pyrimidine 5’-nucleotidase (PYR5), Protein tyrosine phosphatase 1 (PTP1), Abscisic aldehyde oxidase 3 (AAO3), ribosomal protein S3a (RPS3a), Ornithine decarboxylase 1 (ODC1), Trehalose-6-phosphate synthase 1 (TPS1), Starch branching enzyme I (SBEI), and Glutathione S-transferase tau 1 (GSTU1), suggesting functional convergence in stress response, metabolism, and development. The SBEI gene, which plays a role in starch metabolism and grain quality, is highlighted in red. **h**, A bar plot showing how much each genome contributes to the pangenome-exclusive orthogroups. Source data are provided in the Source Data file.

### Structural Variants Underpin Genomic Diversity and Adaptive Traits in Elite ex-PVP Sorghum Lines

A total of 1,118,641 SVs were identified across the pangenome, including insertions, deletions, duplications, inversions, and translocations (Fig. 3a), with the highest individual SVs (42,096) found in wild haplotype 1 (Supplementary Table 6). Insertions and deletions had median sizes of 319 bp and 346 bp, respectively, reflecting a predominance of short to medium-sized INDELs with potential regulatory or coding impacts (Fig. 3b). SVs also overlapped promoter and exon regions, with mean lengths of 1,744 bp and 2,000 bp, respectively (Fig. 3c), suggesting functional consequences for gene expression and/or protein activity. Macro-synteny analysis revealed broadly conserved chromosomal architecture with lineage-specific large-scale rearrangements, including a ~5 Mb inversion on chromosome 5 annotated to a wall-associated kinase (WAK) gene associated with cold tolerance (Fig. 3d). Pairwise comparisons across the pangenome highlighted extensive SV-based divergence, particularly between wild and elite ex-PVP accessions (Fig. 3e, Supplementary Table 7). SVs were unevenly distributed across the genome, with higher densities in pericentromeric and telomeric regions, consistent with suppressed recombination and TE accumulation (Fig. 3f) ^51^.

**Fig. 3.**
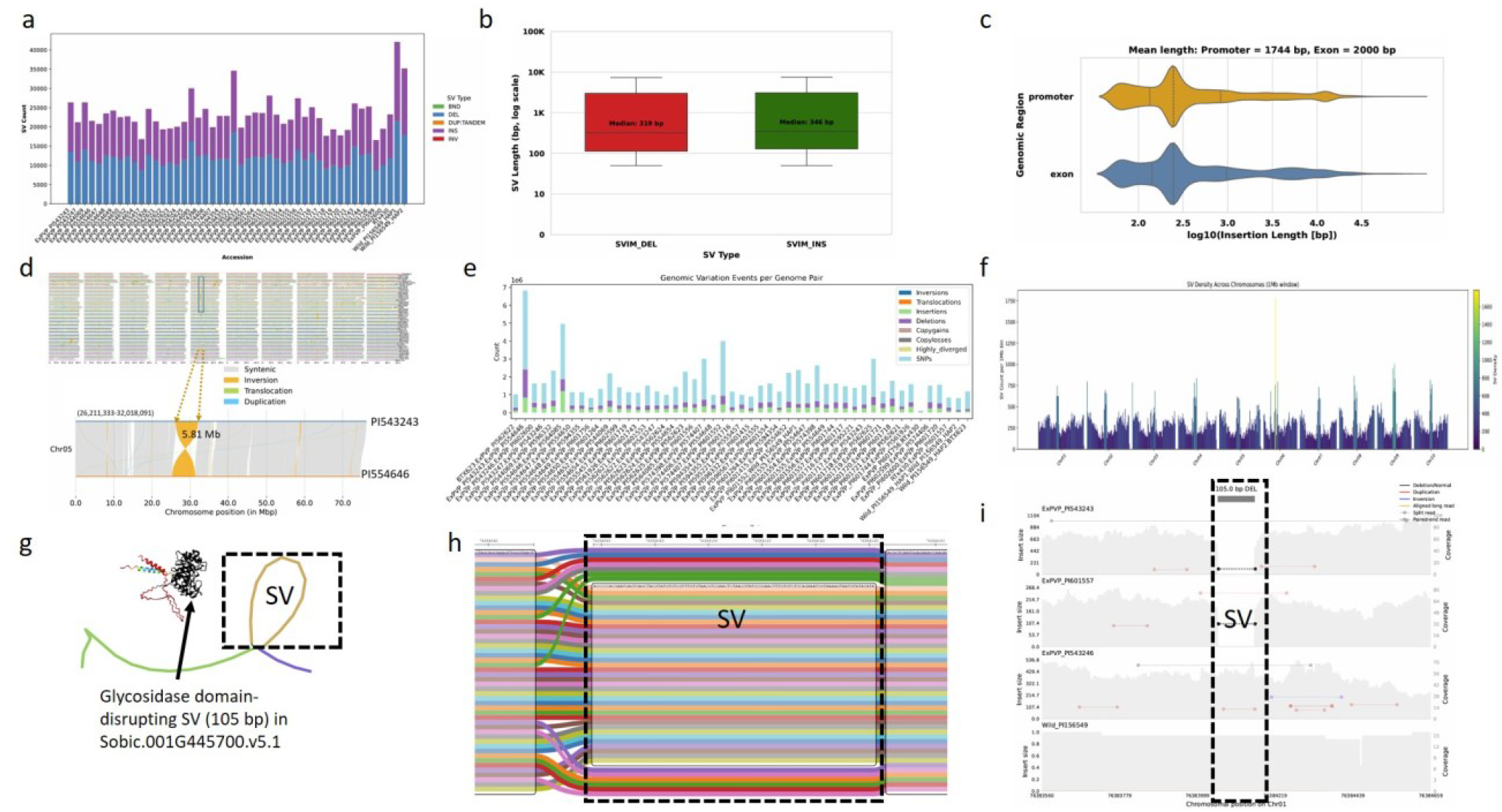
Structural Variants in the Sorghum Pangenome Impact Gene Function and Stress-Related Pathways. **a**, Total number of structural variants (SVs) identified across the 50-genome pangenome; wild haplotype 1 exhibited the highest SV count (42,096), highlighting elevated genomic diversity. **b**, Length distribution of insertions and deletions (INDELs), highlighting the prevalence of short to medium-sized variants. **c**, Distribution of SVs lengths within promoter and exon regions, with potential impacts on gene regulation and coding sequences. **d**, Macro-synteny karyotype plot illustrating conserved chromosomal structure and large-scale rearrangements across sorghum genomes, including a ~5 Mb inversion on chromosome 5 linked to a wall-associated kinase (WAK) candidate gene associated with cold tolerance. **e**, Distribution of pairwise genomic differences across the pangenome, illustrating inter-accession genomic diversity. **f**, Genome-wide SVs density along sorghum chromosomes, with enriched regions corresponding to pericentromeric and telomeric zones. **g**, Pangenome graph and 3D structural visualization of the *Sobic.001G445700.v5.1* protein, a homolog of the *Arabidopsis thaliana β-1,3-GLUCANASE A6* gene involved in plant defense and cell wall remodeling. A 105 bp deletion in this gene disrupts the conserved glycosidase domain (black), likely impairing enzymatic function and suggesting a potential breeding signature for reduced defense investment under high-input agricultural systems. **h**, Presence-absence pattern of the *Sobic.001G445700.v5.1* SVs across the pangenome, distinguishing elite lines from wild accessions. **i**, In silico validation of the structural variant, based on whole-genome read alignments viewed in IGV, confirmed genotype-specific loss of functional sequence—observed in accessions with the 105 bp deletion (ExPVP_PI543243, ExPVP_PI601557) compared to those without (ExPVP_PI543246, Wild_PI565549). Source data are provided in the Source Data file.

Among key SVs, a 105 bp deletion was detected in *Sobic.001G445700.v5.1*, a gene homologous to *Arabidopsis thaliana β-1,3-GLUCANASE A6* gene. The deletion removes a conserved glycosidase domain (Fig. 3g), likely impairing its role in cell wall remodeling and defense. The variant is predominantly present in elite ex-PVP lines and absent in wild accessions, which retain the full gene (Fig. 3h), suggesting this deletion may reflect relaxed defense investment under high-input agricultural systems (e.g., pesticide applications). This pattern could result from reduced pathogen pressure in managed environments or represent a trade-off favoring growth and yield over defense. Whole-genome alignments further confirmed genotype-specific loss of functional sequence, as evidenced by the absence of mapped reads in lines carrying the deletion (Fig. 3i). These findings highlight SVs as a distinct and functionally relevant layer of genomic variation contributing to cultivated sorghum diversity and productivity.

### Selection Targets Circadian and Photoperiod Genes in U.S. Sorghum

We aligned the whole genome sequences of 46 ex-PVP and 25 wild sorghum accessions (Supplementary Table 1) to the ‘BTx623’ reference genome. SNPs were called across all accessions, and after quality filtering, 34,035 high-confidence biallelic SNPs were retained for population structure analysis. Using this dataset, ADMIXTURE analysis (K = 1–10) identified K = 2 as the optimal number of ancestral populations based on the lowest cross-validation error (Supplementary Fig. 2a-b). PCA on the same SNP dataset further detected three genetic clusters, separating ex-PVP lines from two distinct wild subgroups (wild1 and wild2; Supplementary Fig. 2c-f). Based on this population structure, we performed genome-wide selection scans using FST, XP-CLR, and π-ratio to identify genomic regions under differential selection between wild and cultivated accessions.

Selective sweep analysis revealed that XP-CLR and π-ratio identified the most candidates, while overlapping regions from all three methods defined smaller, high-confidence gene sets under strong selection (Supplementary Tables 8-9). Multiple sweep regions differentiated wild and elite ex-PVP lines (Fig. 4a, Supplementary Fig. 3b-d), with strong signals co-localizing with known domestication genes involved in auxin transport, seed dormancy, and aromatic amino acid biosynthesis (Fig. 4b). These pathways are critical for traits such as plant architecture, seed dispersal, and germination.

**Fig. 4.**
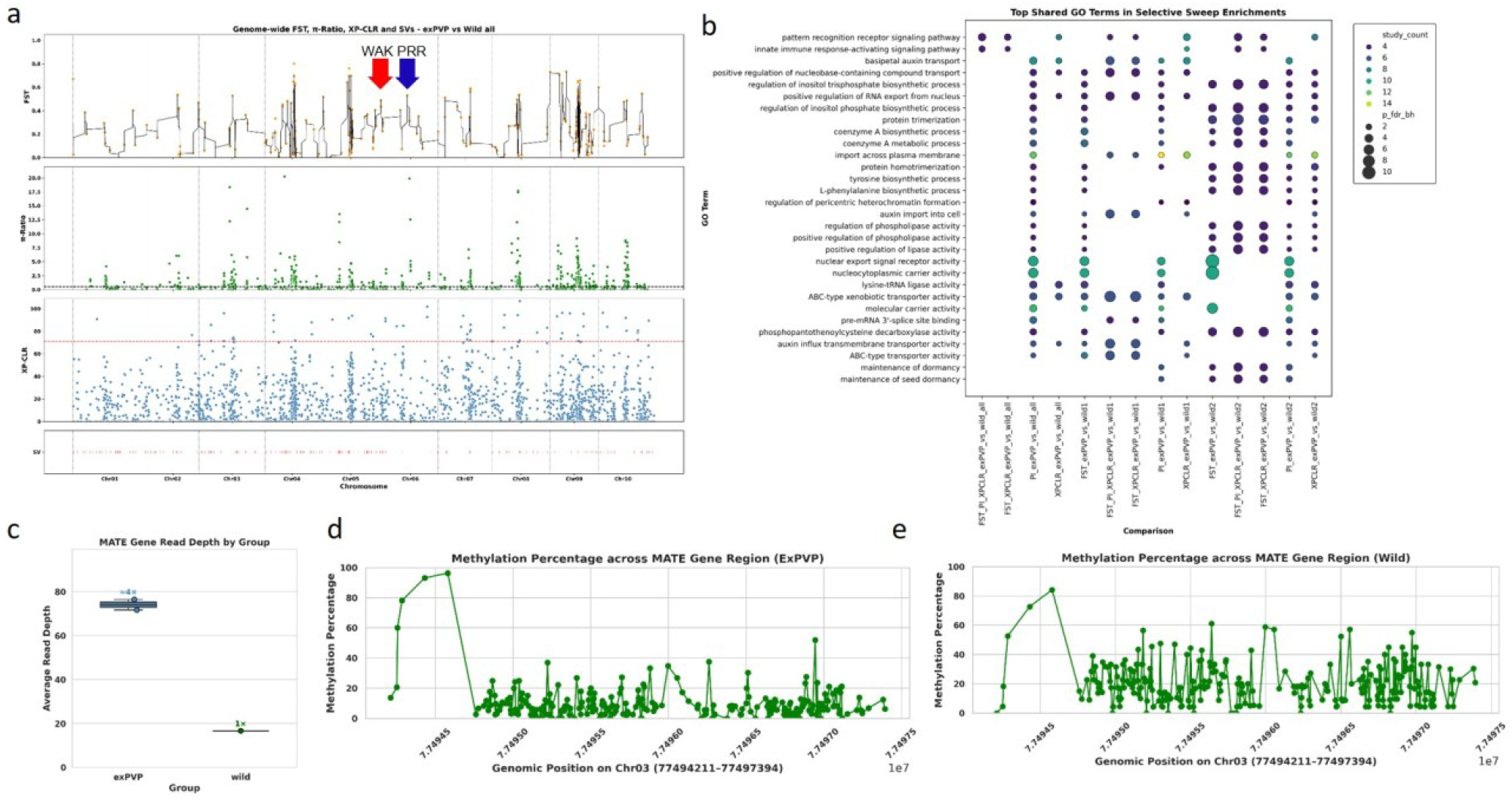
Selection Signatures During Sorghum Improvement Reveal Functional Divergence and Adaptive Loci. **a**, Genome-wide selective sweep signals (FST, π, XP-CLR) across chromosomes comparing 25 wild vs. 46 ex-PVP accessions. The bottom track shows large structural variants identified across the chromosomes. The red arrow marks the position of a *WALL-ASSOCIATED RECEPTOR KINASE* gene *(Sobic.005G179200*) on chromosome 5 (~71 Mb), a member of a gene family known to regulate cell wall integrity and stress signaling under cold conditions. The blue arrow indicates the *PSEUDO-RESPONSE REGULATOR 37* gene (*SbPRR37, Sobic.006G057866*) on chromosome 6 (~41 Mb), a key regulator of photoperiod sensitivity and flowering time in sorghum. **b**, Gene Ontology (GO) enrichment of genes within top 5% sweep regions, highlighting domestication and improvement functions (e.g., auxin transport, immune signaling, dormancy) from various population comparisons and methods shown on the x-axis. **c**, Copy number variation of the *MULTIDRUG AND TOXIC COMPOUND EXTRUSION (MATE*) gene between representative wild and ex-PVP lines, showing approximately fourfold higher copy number in the ex-PVP lines. **d**, methylation percentage across *MATE* gene loci in ex-PVP, indicating low levels across the gene body region. **e**, methylation percentage across *MATE* loci in wild accession showing dynamic levels across the gene body region. Both accessions exhibit higher methylation levels in the promoter region. Source data are provided in the Source Data file.

Although a *β-1,3-GLUCANASE* showed structural disruption in ex-PVP lines (Fig. 3g-i), genome-wide sweeps reveal positive selection on other immune-related pathways, such as *PATTERN RECOGNITION RECEPTOR (PRRs)* signaling and innate immune activation, suggesting a shift in defense strategy rather than an overall reduction. Enriched GO terms also indicate that modern breeding emphasized immune signaling, xenobiotic transport, phospholipase activity, and RNA export (Fig. 4b)—reflecting a broader focus on stress resilience and metabolic efficiency under high-input agricultural systems.

Enrichment analysis of genes under selection highlighted key regulators of circadian rhythm and flowering time, including *PRR7* and *PRR95*, and *PRR95*, core components of the circadian clock, as well as *CO* and *PHOT1*, which mediate photoperiodic flowering (Supplementary Table 10). Genes involved in seasonal development—such as *FRIGIDA (FRI), VRN2B*, and *VRN3B*— were also enriched, suggesting coordinated selection on flowering pathways. Additional targets included *REGULATORY-ASSOCIATED PROTEIN OF TOR 1A (RAPTORA), TARGET OF RAPAMYCIN (TOR), GRAIN NUMBER, PLANT HEIGHT, AND HEADING DATE 7 / MATURITY 6 (GHD7/MA6)*, and *XAP5 CIRCADIAN TIMEKEEPER (XCT)*, implicating broader roles for metabolic and growth regulation in adaptation ^52^. These loci lie within sweep regions analogous to those reported in other crops—for example, selection on *DRIVERS OF THE CIRCADIAN CLOCK (DOC)* in barley has been shown to modulate rhythmicity and reproductive traits under domestication ^53^. The co-selection of these interconnected regulators underscores a multifaceted strategy favoring clock-modulated traits, offering breeders a powerful entry point to fine-tune flowering time, growth rate, and environmental responsiveness in modern sorghum improvement programs.

Further analysis of genomic regions associated with flowering time, photoperiod sensitivity, and circadian regulation revealed a strong signal at the *SbPRR37 locus (Sobic.006G057866)*, the classical *MATURITY 1 (Ma1)* gene in sorghum. Its closest *Arabidopsis thaliana* ortholog is *PRR7* (Fig. 5a, Supplementary Table 11). We detected CNVs at *SbPRR37*, with two copies present in most cultivated genotypes but only one in wild haplotype 2, suggesting reduced redundancy or differential selection pressure in wild populations. Additional sweep intervals contained other circadian and flowering regulators, including *PRR7* (Sobic.001G411400), PRR95 (*Sobic.002G275100*), *Ma2* (*Sobic.002G302700*), and *GDH7/Ma6* (*Sobic.006G004400*), supporting the hypothesis that multiple maturity genes (*Ma1–Ma6*) have been targets of selection during domestication and improvement ^37^.

**Figure 5.**
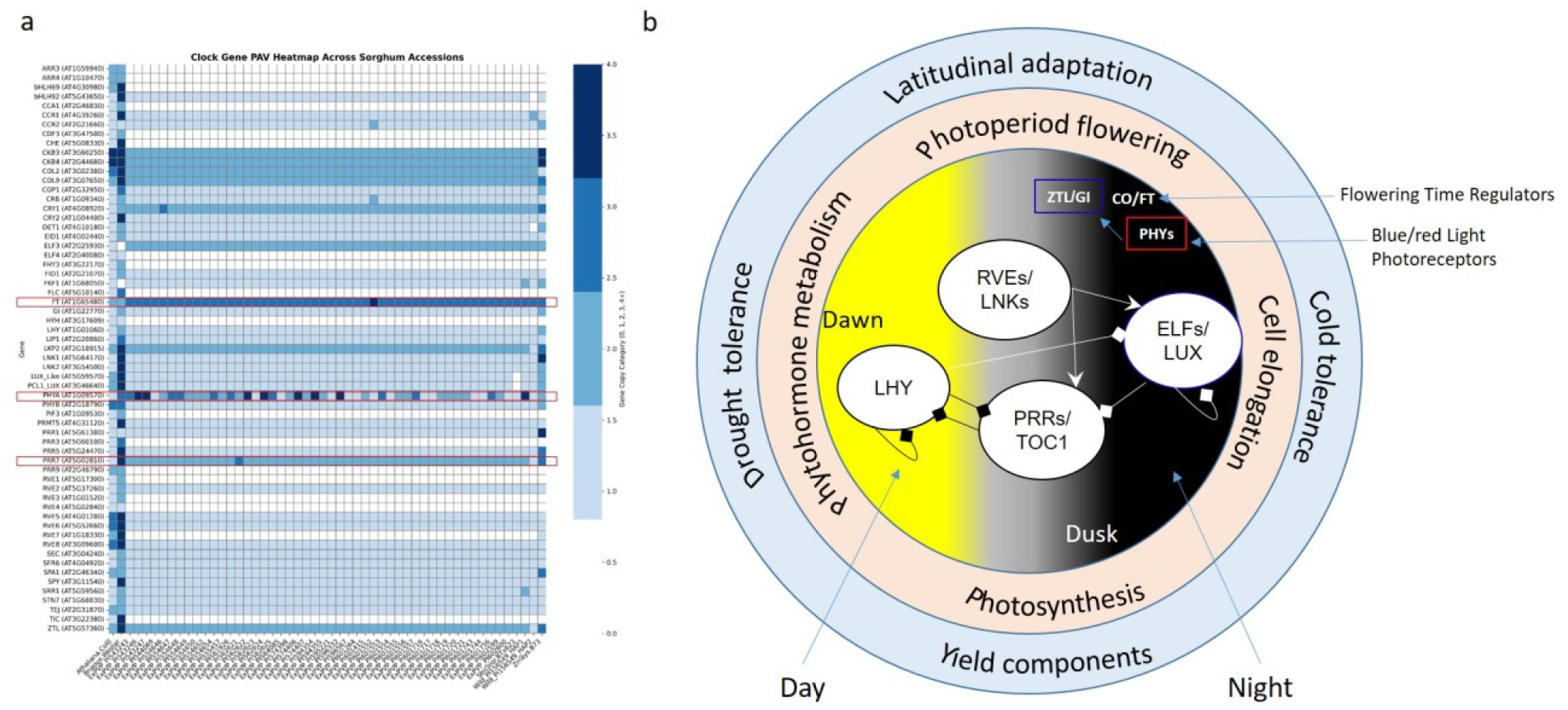
Time-of-Day (TOD) Gene Variation and Model of the Sorghum Circadian Clock Network. **a**, Variation in circadian clock and flowering-time genes reveals distinct selection pressures in cultivated sorghum. Copy number expansion of the circadian photoreceptor PHYTOCHROME A (PHYA) and the clock regulator PSEUDO-RESPONSE REGULATOR 37 (SbPRR37, Sobic.006G057866)—orthologous to PRR7 (AT5G02810) and PRR3 (AT5G60100) in Arabidopsis thaliana—reflects selection on the circadian system, while duplication of the florigen gene FLOWERING LOCUS T (FT) indicates selection on flowering-time pathways. PHYA and FT are highlighted in red boxes. These patterns suggest targeted adaptation to seasonal and latitudinal environments via independent tuning of clock and flowering components. **b**, Schematic representation of interlocked transcriptional feedback loops that drive circadian rhythms, allowing coordination with environmental signals such as light and temperature. The central area illustrates core clock regulatory interactions, adapted from Bendix et al. (2015) ^37^. Yellow and black regions represent day and night, respectively. Arrows with pointed ends indicate activation; diamond-tipped arrows indicate repression. The orange-shaded middle layer denotes signaling and physiological pathways directly regulated by the circadian clock, while the outer blue-shaded ring highlights traits influenced by clock-controlled outputs. See supplementary notes for details. Source data are provided in the Source Data file.

Several other sweep regions harbored genes involved in florigen signaling and circadian entrainment, such as *VERNALIZATION 3B (VRN3B, Sobic.003G173032*), *CONSTANS (CO, Sobic.010G115800*), and *PHOTOTROPIN 1 (PHOT1, Sobic.008G001000*). Selection signals also overlapped with key metabolic and photomorphogenic regulators, including *TARGET OF RAPAMYCIN (TOR)* (Sobic.009G109200), *REGULATORY-ASSOCIATED PROTEIN OF TOR 1A (*RAPTORa) (Sobic.005G008800), *XAP5 CIRCADIAN TIMEKEEPER (XCT)* (Sobic.002G277600), and *DE-ETIOLATED 1 (DET1)* (Sobic.007G058100). Structural variation, including PAVs and CNVs, was also evident at the *PHYTOCHROME A (PHYA)* and *FLOWERING LOCUS T (*FT), alongside enrichment of core circadian components. Notably, regulators such as *CIRCADIAN CLOCK ASSOCIATED 1 (CCA1)* appeared to be dicot-specific—present in *Arabidopsis thaliana* and *Brassica napus* but absent from the sorghum genome (Fig. 5).

### Copy Number Expansion and Differential Methylation of the MATE gene in ExPVP lines

CNVs analysis based on read depth revealed four copies of the *MATE* (*Sobic.003G403000.v5.1*) gene in ex-PVP lines compared to a single copy in the wild accession (Fig. 4c). This copy number expansion coincides with a marked reduction in DNA methylation across the gene body in the ex-PVP lines, whereas the wild accession exhibited higher and more variable methylation across the same region (Fig. 4d). The lower methylation in ex-PVP accessions, particularly within the gene body (Supplementary Fig. 4), suggests epigenetic reprogramming likely associated with transcriptional activation of duplicated *MATE* copies. In contrast, the higher and fluctuating methylation levels in the wild accession may reflect transcriptional repression or tighter regulation (Fig. 4e). These patterns are consistent with selective amplification and epigenetic remodeling of *MATE* in elite germplasm to support traits such as xenobiotic transport or abiotic stress response ^54^.

## Discussion

This study explores SVs, gene PAVs, and epigenomic remodeling in sorghum, uncovering previously unrecognized diversity linked to key agronomic traits and environmental adaptation, a variation often missed when relying on a single reference genome ^1,21,47,55^.

A substantial fraction of TEs in the pangenome remained unclassified, likely reflecting lineage-specific or divergent TEs families underrepresented in existing repeat libraries ^45,46^. These findings are consistent with recent large-scale pangenome efforts in sorghum, where more than 200 Mb of novel sequence were uncovered across cultivated and wild accessions ^21,22,56^. Our analysis identified 28% of genes displaying PAVs, highlighting genomic diversity among elite lines. This proportion is comparable to elite maize accessions, where ~32% of gene transcripts in elite inbred lines display PAVs and are linked to stress-and environment-responsive functions as well as heterosis ^57^.

The PAVs suggest that multiple gene combinations may have been retained or selected to optimize adaptation and productivity across diverse growing environments. We discovered *SBE1* gene among pangenome-exclusive genes. In cereals like maize and rice, *SBE1* is associated with grain digestibility and quality, suggesting that its presence in elite U.S. sorghum likely reflects selection for processing traits and end-use preferences ^58^.

Despite the pangenome showing strong genome-wide collinearity and preserved telomere architecture, consistent with counter-selection against large-scale rearrangements ^59^, we identified a notable ~5 Mb inversion on chromosome 5 in ex-PVP lines. This region includes a WAK gene, part of a family known to regulate cell wall integrity and stress signaling in cold, drought, and salinity conditions ^60,61^. In rice and Arabidopsis, WAK genes play central roles in early abiotic stress responses and cell wall remodeling, suggesting that this inversion may represent targeted selection for cold tolerance in temperate breeding programs.

Our genome-wide selective sweep analysis revealed strong signals of selection on circadian and photoperiodic regulators, extending beyond canonical domestication genes ^31^. We detected a sweep at *SbPRR37 (Sobic.006G057866)*, the gene underlying the *Ma1* locus, a major repressor of flowering under long-day (LD) conditions. Recessive *ma1* alleles historically enabled early flowering and adaptation to temperate latitudes ^62,63^, while dominant *Ma1* alleles have been favored in modern biomass sorghum lines for delayed flowering under LD conditions by increasing expression of *SbCO* which repress *SbFT* thus extended vegetative growth ^64,65^. Our observation of CNVs at *SbPRR37*—with two copies in most cultivated lines and one in wild haplotype 2—suggests differential selective pressure on photoperiod response across gene pools.

Additional sweeps at *PRR7, PRR95, Ma2, Ma6, VRN3b*, and *CO* underscore the polygenic nature of flowering-time control, integrating both classical *Ma* loci and newly implicated regulators. Notably, genes such as *PHYA* exhibited copy number expansion in ex-PVP lines, which may enhance photoperiod sensitivity or stabilize developmental transitions under variable seasonal cues. The detection of enriched selection signals at *PHOT1, DET1*, and *XAP5* suggests that upstream regulatory pathways have been knowingly or unknowingly targeted during sorghum improvement. This highlights the importance of fine-tuned seasonal and environmental sensing in shaping phenological plasticity. Similar patterns have been observed in barley, where a natural variation at *Ppd-H1*, a *PRR7* ortholog, alters the circadian expression of flowering time genes without disrupting core clock components, enabling latitude-specific flowering responses ^66^. In maize, selection on *ZmCCT9*, a diurnally regulated repressor of the florigen *ZCN8*, delays flowering under LD conditions and enhances adaptation to higher latitudes ^67^.

Beyond structural and genetic variation, we observed epigenomic divergence at functionally relevant loci, consistent with evidence that epigenetic modifications can be subject to selection and contribute to rapid adaptation, as demonstrated in *Arabidopsis thaliana* ^*68*^. Notably, MATE transporter genes exhibited both a fourfold copy number expansion and reduced gene-body methylation in ex-PVP lines. These transporters mediate the efflux of organic acids, xenobiotics, and flavonoids, with established roles in herbicide detoxification and aluminum tolerance in cereals ^69–72^. The observed gene-body hypomethylation may enhance expression under high-input agricultural conditions, suggesting that coordinated structural and epigenetic remodeling has been a feature of modern sorghum breeding.

Together, these results support a model in which sorghum’s adaptation to temperate climates and agricultural intensification has been shaped by diverse molecular mechanisms—including targeted selection on flowering-time genes, clock genes, and epigenetic reprogramming of stress and detoxification genes. Our study complements and extends recent sorghum pangenome analyses ^21,22,56,73^, which established foundational catalogs of structural and gene-content diversity across races and wild sorghum relatives. While those efforts focused on broad germplasm panels, our study focuses on ex-PVP cultivars—historically important but under-characterized—and adds critical insights into how modern breeding has reshaped the genome.

Although this study offers major insights, limitations remain. The ex-PVP panel studied here represents lines from 1976–1992 and may not reflect recent commercial diversity. The functional impacts of structural variants and epialleles also require validation. Future work should integrate long-read resequencing, qPCR, and CRISPR to resolve complex loci. Additionally, field-based time-series transcriptomics are needed to link genomic variation with gene expression under natural conditions, improving our understanding of trait-environment interactions and guiding precision breeding.

In conclusion, this study demonstrates how modern breeding has shaped the sorghum genome through targeted structural, regulatory, and epigenetic changes, offering new opportunities for trait dissection and breeding innovation. The resulting pangenome resource enhances our ability to explore adaptation, gene-trait relationships, and the development of resilient, climate-smart sorghum varieties to meet global food, feed, and bioenergy demands.

## Methods

### Plant Materials

Seeds for 46 elite ex-Plant Variety Protection (ex-PVP) sorghum lines were obtained from the USDA Germplasm Resources Information Network (GRIN). These accessions originated from the breeding programs of Pioneer Hi-Bred International, Inc. (n = 39), Novartis Seeds, Inc. (n = 1), Cargill Wheat Research Farm (n = 1), Holden’s Foundation Seeds, Inc. (n = 1), Walter Moss Seed Company, LLC (n = 1), Ring Around Products, Inc. (n = 1), and Northrup, King & Company (n = 2). Wild sorghum accessions were also sourced from GRIN and selected to represent diverse geographic and taxonomic origins for comparative genomic analyses. RNA-seq was performed on two sorghum accessions, ‘PI329478’ and ‘PI510757’, using eight tissues per genotype: seedling, 3-leaf, 5-leaf, tiller, boot, panicle with anthers, root, and dough-stage panicle. Total RNA was extracted, quality-checked, and used to construct poly(A)-selected libraries, which were sequenced on an Illumina platform to produce paired-end reads (Supplementary Table 12).

### High-Molecular-Weight (HMW) DNA Extraction and Sequencing

HMW genomic DNA was extracted from 3-week-old seedlings using Carlson lysis buffer and Qiagen Genomic-tips, following the Oxford Nanopore Technologies (ONT) ‘Plant leaf gDNA’ protocol for Arabidopsis. Samples were ground in liquid nitrogen, followed by two chloroform:isoamyl alcohol (24:1) washes. Total Pure NGS beads (Omega Bio-tek) were used for cleanup. DNA was size-selected for fragments >10–25 kb using the ONT Short Fragment Eliminator Kit (EXP-SFE001) and quality assessed using the Agilent Genomic DNA ScreenTape system (5067-5365) or Femto Pulse Genomic DNA 165 kb Kit (FP-1002-0275). HiFi circular consensus sequencing (CCS) libraries were prepared using the PacBio protocol (PN 101-853-100 v5). Genomic DNA was sheared using Covaris g-TUBEs to a modal fragment size of ~18 kb, and fragments <10 kb were excluded using BluePippin S1 High Pass 6–10 kb cassettes.

### Genome Assembly and Annotation

Ex-PVP genomes were assembled from Oxford Nanopore Technologies (ONT) reads using Flye (v2.9.2), followed by polishing with Racon (v1.5.0) and Pilon (v1.24) using Illumina short reads. Wild genomes were assembled using Hifiasm (v0.16.1-r159) with PacBio HiFi reads. Assemblies were scaffolded using RagTag (v2.1.0) with the BTx623 reference genome ^16,74^. For the wild accession ‘PI156549’, Hi-C reads were additionally aligned to contigs using the Juicer pipeline (v1.6.2), followed by scaffolding with 3D-DNA (v180419). Manual curation was performed using Juicebox Assembly Tools (v1.11.08). Genome assembly quality was evaluated using BUSCO (v5.4.3) with the embryophyta_odb10 dataset and the LTR Assembly Index (LAI), which provides a metric for assembly continuity and repeat completeness based on intact long terminal repeat retrotransposons. Gene models were predicted using Helixer, a deep learning-based annotation tool trained on plant genomes ^44^.

### Transposable Elements (TEs) Detection

TEs were annotated using EDTA (v2.0.0) with the flags --sensitive 1 and --anno 1, which integrates both structure- and homology-based approaches. The pipeline incorporates multiple tools, including LTRharvest, LTR_FINDER, LTR_retriever, Generic Repeat Finder, TIR-learner, HelitronScanner, TEsorter, RepeatModeler, and RepeatMasker. Interspersed and tandem repeats were further annotated using RepeatMasker (v4.1.1) with the custom EDTA-derived TE library, and tandem repeats were identified with TRF (v4.09).

### Large Structural Variant Detection

Structural variants (SVs) were identified using a combination of reference-based and assembly-based approaches based on 46 exPVP and wild accession PI156549. SYRI (https://github.com/schneebergerlab/syri) was first used to characterize large-scale structural rearrangements, including translocations and inversions. For assembly-based SV calling, Svim-asm (v1.0.3) was applied to genome assemblies to detect insertions, deletions, and complex events. Additionally, CuteSV (v2.1.2) was used to extract duplications (DUP), inversions (INV), and breakends (BND). SV results were refined and benchmarked using Truvari (v5.3.0), and nonredundant SV sets were merged using SURVIVOR (v1.0.7). Whole-genome alignments were performed using Minimap2 (v2.29-r1283) to support SV analysis. Pangenome graph construction and structural diversity visualization were performed using PGGB (v0.6.0).

### Synteny and Collinearity Analysis

Conducted at both genome and gene levels using D-Genies (v1.4) with whole-genome alignments from Minimap2 (v2.22), and MCScan in the JCVI package (v1.2.7) based on CDS alignments of longest isoforms generated using Last (v1418).

### Identification of Orthologs

The interspecific all orthogroup gene pairs were determined using OrthoFinder (v2.5.4, ortho_opt: -M msa -A mafft -T fasttree)

### Single-Nucleotide Variant Calling, Admixture Analysis, and Population Structure

Genome assemblies from 46 ex-PVP lines and 25 wild sorghum accessions (Supplementary Table 1) were aligned to the BTx623 v5 reference genome ^16,75^, and variants were called using FreeBayes (v1.3.6) with a ploidy-aware setting (−g 1000). Variants were filtered to retain sites with quality score (QUAL) > 30, yielding 494,424 SNPs and 33,132 INDELs (Source: Fig4_Sbic2groups_wild_cultivated_variants.v1.freebayes.filt.vcf.gz). Filtered SNPs were further refined and summarized using vcftools (v0.1.16) and rtg tools (v3.12) to retain only biallelic sites with MAF between 0.01 and 0.99 and no more than 10% missing data, resulting in 34,035 high-quality SNPs. These were used for population structure analysis. ADMIXTURE (v1.3.0) was run with five-fold cross-validation for K = 1 to 10, identifying K = 2 as the optimal number of ancestral populations based on the lowest CV error (Supplementary Fig. 2a-b). Population structure was also assessed using principal component analysis (PCA) in PLINK (v1.90b7.7). PCA separated accessions into three distinct clusters, with PC1 and PC2 explaining 30.34% and 11.42% of the variation, respectively (Supplementary Fig. 2c-f).

### Selection Signature Identification

We performed four pairwise comparisons using filtered SNPs derived from whole-genome sequencing data: #1. Ex-PVP lines vs. all wild accessions. #2. Ex-PVP lines vs. Wild1. #3. Ex-PVP lines vs. Wild2. #4. Wild1 vs. Wild2. For each comparison, we employed three complementary methods to detect selective sweeps: FST (fixation index):

Population differentiation was estimated using the Weir and Cockerham FST method implemented in VCFtools. Analyses were performed using sliding windows of 100 kb with 10 kb steps (−-fst-window-size 100000 --fst-window-step 10000). Windows with FST > 0.3 were considered indicative of strong differentiation. π-ratio (nucleotide diversity ratio): Nucleotide diversity (π) was computed separately for each group using VCFtools (−-window-pi 100000 -- window-pi-step 10000), and the ratio π exPVP / π wild was calculated. Windows with a π-ratio < 0.5 were interpreted as signals of reduced genetic diversity in the ex-PVP group, consistent with selection. XP-CLR (Cross-Population Composite Likelihood Ratio): XP-CLR scores were computed using XP-CLR (v1.1.2, https://github.com/hardingnj/xpclr), using a window size of 100 kb and step size of 10 kb. Windows in the top 1% (99th percentile) of XP-CLR scores were considered candidate regions under strong selection. We further defined high-confidence sweep regions using FST & XP-CLR intersection and FST & π-ratio & XP-CLR intersection. Only windows meeting all three thresholds were retained in the final sweep set, enhancing confidence that these represent genuine targets of selection during domestication or improvement.

### DNA Methylation Analysis

LoReMe (Long Read Methylation) (Oxford Nanopore Technologies ·GitHub) was used to infer DNA methylation patterns. In brief, the process involved the conversion of ONT FAST5 data to POD5 format using the Loreme Dorado-convert tool (v.0.3.1). Subsequently, super-high-accuracy base calling was performed, aligning the sequences to the reference genome. Modkit (v0.1.11) was employed to generate a bed file containing comprehensive methylation data, enabling us to create visual representations of methylation profiles for further investigation and interpretation.

### Circadian Clock, Photoperiod, and Flowering Time Genes

Sorghum core circadian clock, light signaling, and flowering time orthologs genes were identified using known *Arabidopsis thaliana* genes as the seed ^76^

### Gene Ontology (GO) Enrichment Analysis

Genes located within or overlapping candidate sweep regions (±100 kb) were extracted by intersecting sweep windows with mRNA coordinates from the Sorghum bicolor BTx623 v5.1 reference annotation (Phytozome^16^). This was done using the PyRanges Python library. Genes from each metric-specific sweep set (e.g., FST-only, XP-CLR-only, triple-intersection) were analyzed separately. GO enrichment analysis was conducted using the GOATOOLS Python library (v1.2.3) with the find_enrichment.py script. GO term associations were generated from functional annotations predicted by eggNOG-mapper, and formatted into id2gos style association files. The GO DAG structure was derived from go-basic.obo. Each gene list was tested against a common background of all BTx623 genes with GO annotations. Statistically significant GO terms (corrected p-value < 0.05, using Benjamini-Hochberg FDR) were retained and reported.

## Data Availability

All relevant datasets are provided in the main text and supplementary materials. Raw sequencing reads are publicly available under NCBI BioProject PRJNA1283509.

Analysis scripts and pipelines are available on GitHub at https://github.com/jkitony/Pangenome-of-U.S.-ex-PVP-and-Wild-Sorghum.

Genome assemblies and annotations can be accessed via the Salk Institute’s resource portal (https://resources.michael.salk.edu/), Sorghumbase (https://www.sorghumbase.org/), and Figshare (https://doi.org/10.6084/m9.figshare.29261795.v4).

## Acknowledgements

We would like to thank the Donald Danforth Plant Science Center, the United States Department of Agriculture (USDA), and the Germplasm Resources Information Network (GRIN) for providing resources used in this study. This work was supported through the Salk Harnessing Plants Initiative (HPI) with funding from the TED Audacious, Bezos Earth Fund, and Hess Corporation.

