## Supplementary Information for "Pangenome of U.S. ex-PVP and Wild Sorghum Reveals Structural Variants and Selective Sweeps Shaping Adaptation and Trait Improvement"

**Running Title:** Pangenome of U.S. ex-PVP and Wild Sorghum

**Authors**
Justine K. Kitony [1], Emily R. Murray [1], Kelly Colt [1], Ryan C. Lynch [1], Nicholas Allsing [1], Nolan T. Hartwick [1], Tiffany Duong [1], Jocelyn Saxton [2], Nadia Shakoor [2], Todd P. Michael [1,3]

**Affiliation**
1. Plant Molecular and Cellular Biology Laboratory, Salk Institute for Biological Studies, La Jolla, CA, USA
2. Donald Danforth Plant Science Center, Olivette, MO, USA
3. Science and Conservation, San Diego Botanical Garden, Encinitas, CA, USA

**Corresponding Author**
Todd P. Michael,

**Key Words**
Plant Variety Protection (ex-PVP), sorghum pangenome, selective sweep, structural variants (SVs), long-read sequencing, chromosome-level assemblies, presence–absence variation (PAVs), and copy number variation (CNVs)

**Supplementary Notes**……………………………………………………………...…… 2-6

**Supplementary Figures**........................................................................................ 7-10

**Supplementary Tables**......................................................................................... 11-19

**Supplementary References**................................................................................. 20-21

### Supplementary Notes

#### Sample Selection

We assembled a panel comprising 46 elite sorghum lines formerly protected under the U.S. Plant Variety Protection (ex-PVP) system, which confers protection for 20 years, together with a set of diverse wild accessions. The ex-PVP lines, registered between 1976 and 1992, were sourced from the USDA Germplasm Resources Information Network (GRIN) and reflect historical commercial breeding efforts by Pioneer Hi-Bred International, Inc. (n = 39), Novartis Seeds, Inc. (n = 1), Cargill Wheat Research Farm (n = 1), Holden's Foundation Seeds, Inc. (n = 1), Walter Moss Seed Company, LLC (n = 1), Ring Around Products, Inc. (n = 1), and Northrup, King & Company (n = 2). Agronomically, the ex-PVP lines represented a broad range of tillering phenotypes, from low-tillering genotypes optimized for grain production to moderately and highly tillering types with potential forage or dual-purpose utility.

The ex-PVP phenotypic diversity underscores their relevance for investigating the genomic underpinnings of sorghum's adaptation to different cropping systems. To enable comparative genomic analyses, we also included wild sorghum accessions from GRIN, selected to maximize both phylogenetic and geographic diversity (Supplementary Fig. 2). Among these, *Sorghum bicolor* subsp. *verticilliflorum* accession ‘PI 156549’, originating from Zimbabwe, was selected for Hi-C scaffolding and inclusion in pangenome construction. This Rhodesian sudangrass type has historically contributed to the development of male-sterile lines with superior forage potential and may represent one of the ancestral contributors to elite hybrid forage sorghums [1,2].

#### Genome Assembly and Annotation

We generated chromosome-scale genome assemblies for 46 ex-PVP lines and several wild Sorghum accessions to support pangenome construction and comparative genomic analyses. Ex-PVP genomes were assembled using long-read ONT data with hybrid polishing, while wild accessions were assembled using PacBio HiFi reads. Scaffolding was reference-guided using RagTag with the ‘BTx623’ reference [3,4]. One wild accession, ‘PI 156549’, was additionally scaffolded with Hi-C data and manually curated, resulting in a high-quality representative genome for the wild group [5].

Assembly quality was assessed using BUSCO (embryophyta_odb10) to evaluate completeness [6], and the LTR Assembly Index (LAI) to assess repeat continuity [7]. Gene prediction was performed using Helixer, providing consistent structural annotations across accessions [8]. Transposable elements (TEs) were annotated using the EDTA pipeline, integrating structure-based and homology-based repeat identification [9]. Custom repeat libraries were used with RepeatMasker to mask interspersed and tandem repeats. These annotations support downstream analyses of gene content variation, structural variants, and repeat dynamics analyses within the sorghum pangenome.

#### Gene Family Inference and Pangenome Stratification

We constructed orthologous gene families across 50 sorghum genomes using OrthoFinder (v2.5.4), leveraging sequence similarity and gene tree inference to group genes into orthogroups. This analysis identified 36,004 gene families, which were classified into four categories based on their distribution across genomes: core (present in ≥49 genomes; 71.5%), soft-core (47–48 genomes; 2.2%), dispensable (7–46 genomes; 16.7%), and private (<7 genomes; 9.6%) (Fig. 2a). These classifications capture a spectrum of gene conservation and variation, with core genes reflecting essential functions, while dispensable and private genes likely represent adaptations to specific breeding objectives or environmental niches.

Gene space dynamics were evaluated by generating collector's curves through random sampling of genome combinations. The core genome curve declined asymptotically, while the pangenome curve plateaued, suggesting saturation of novel gene families with the inclusion of additional accessions (Fig. 2b). A power law model of novel gene family discovery ($a = 4.13$, $R^2 = 0.999$) confirmed the closed nature of the gene-based sorghum pangenome (Fig. 2c). This contrasts with the K-mer-based analysis (Supplementary Fig. 1e), which indicated ongoing K-mer accumulation, reflecting structural and non-genic variation not captured at the gene family level.

Comparison to the 'BTx623' reference genome revealed 6,389 orthogroups absent from the reference but present in the broader pangenome (Fig. 2d). These include gene families with annotated KEGG orthologs involved in stress response, such as GLUTATHIONE S-TRANSFERASE TAU 1 (GSTU1) and ABSCISIC ALDEHYDE OXIDASE 3 (AAO3); specialized metabolism, such as FLAVONOID 3'-HYDROXYLASE (F3'H), SALICYLIC ACID CARBOXYL METHYLTRANSFERASE (SAMT), and TREHALOSE-6-PHOSPHATE SYNTHASE 1 (TPS1); energy metabolism, such as ATP SYNTHASE SUBUNIT BETA (ATPB), NADH DEHYDROGENASE SUBUNIT H (NDHH), and NADH DEHYDROGENASE SUBUNIT F (NDHF); and carbohydrate biosynthesis, such as STARCH BRANCHING ENZYME I (SBE1)—many of which may have been selected during breeding for grain quality or digestibility traits.

Gene presence–absence patterns showed consistent clustering among ex-PVP lines (Fig. 2e), and a genome-wide genespace visualization confirmed conservation of genic regions across accessions, with occasional lineage-specific rearrangements (Fig. 2f). These findings demonstrate how gene-based pangenomics complements reference-guided analyses by uncovering lineage-specific diversity and capturing functional variation shaped by breeding and domestication.

**Structural Variant (SVs) Detection and Synteny Analysis**

SVs represent a critical yet underexplored layer of genomic variation in crop species, where reliance on single reference genomes obscures much of the intraspecific diversity shaped by domestication, environmental adaptation, and breeding [10]. Advances in long-read sequencing and multi-assembly pangenomics now enable the detection of SVs at nucleotide to megabase scale, revealing their functional importance in agronomic traits such as yield, defense, and stress response [11]. In this study, we leveraged a haplotype-resolved pangenome comprising 46 elite U.S. ex-PVP lines and one wild accession (PI156549) to systematically characterize SVs using both reference-guided and de novo assembly-based approaches. This framework allowed us to resolve a broad spectrum of SVs, ranging from short INDELs to large chromosomal rearrangements that alter gene collinearity and synteny.

SVs were detected using two complementary workflows:

1. Assembly-based SV Calling:
   Long-read genome assemblies were aligned using Minimap2 (v2.29-r1283), and structural variants were identified with Svim-asm (v1.0.3), which is optimized for high-contiguity assemblies and can resolve complex SVs including insertions, deletions,

and translocations. Additionally, CuteSV (v2.1.2) was employed to extract specific SV types such as duplications (DUP), inversions (INV), and breakends (BND), expanding the scope of detection to include more complex rearrangements.

2. Reference-guided Detection:
   Whole-genome alignments were input to SYRI (https://github.com/schneebergerlab/syri), a tool designed to identify large-scale chromosomal rearrangements, including inversions, translocations, and duplications, by comparing each accession to the reference genome BTx623. This reference-based perspective complements the assembly-based approach by capturing lineage-specific differences in genome organization.

SVIM-asm and CuteSV outputs were filtered and benchmarked using Truvari (v5.3.0) to increase confidence in structural variant (SV) calls and reduce redundancy. High-confidence SVs were subsequently merged with SURVIVOR (v1.0.7), generating a unified SV catalog across the dataset. This workflow minimized tool-specific inconsistencies and improved sensitivity to both shared and accession-specific variants.

A graph-based pangenome was constructed using PGGB (v0.6.0), enabling visualization of the structural landscape and exploration of SV breakpoints in the context of sequence continuity. This graph representation allowed for the detection of allelic variation and sequence-specific loss or gain across the population, particularly for loci implicated in defense or stress response, such as a 105 bp deletion in the *GLUCAN ENDO-1,3-β-GLUCOSIDASE A6* gene (*Sobic.001G445700*), which disrupts a conserved domain and is enriched in elite lines but absent in wild accessions.

In parallel, we explored genome collinearity and synteny conservation at both the chromosome and gene levels. Whole-genome synteny was assessed using D-GENIES (v1.4) with Minimap2 alignments (v2.22), revealing largely conserved macro-synteny punctuated by lineage-specific inversions and structural breaks. For gene-level analysis, MCScan from the JCVI toolkit (v1.2.7) was applied to CDS alignments based on the longest isoforms, generated using LAST (v1418). This approach enabled high-resolution mapping of orthologous gene blocks and allowed us to distinguish conserved versus rearranged segments.

Together, these complementary pipelines revealed that structural variants are pervasive across the sorghum pangenome, with functional implications ranging from altered gene dosage and regulatory landscapes to potential loss of defense genes under relaxed selection in modern breeding. This structural layer adds crucial context to SNP-based and gene presence/absence analyses, underscoring the value of graph-based and multi-genome frameworks in capturing hidden genomic diversity.

**Population Structure and Selection Signatures**

We first performed variant discovery using long-read genome alignments against the ‘BTx623’ v5 reference. Variant calling with FreeBayes (v1.3.6) yielded ~500k raw SNPs and INDELs. After stringent filtering for biallelic SNPs with high confidence ($QUAL > 30$), minor allele frequency ($0.01 \leq MAF \leq 0.99$), and $\leq 10\%$ missing data, we retained 34,035 high-quality SNPs suitable for population-level inference.

Population structure was assessed using two complementary approaches:

- ADMIXTURE analysis (v1.3.0) was performed with cross-validation for K = 1–10, identifying K = 2 as the optimal number of ancestral populations. This partitioning revealed a deep divergence between wild and cultivated accessions, consistent with strong genetic bottlenecks during domestication and improvement.
- Principal Component Analysis (PCA) using PLINK (v1.90b7.7) further separated the 71 accessions into three distinct clusters: one representing the ex-PVP lines and two subgroups within the wild accessions (wild1 and wild2). PC1 and PC2 together explained over 40% of the total variation, highlighting the major axes of sorghum diversification.

To identify genomic regions under selection, we applied three independent but complementary metrics across four pairwise population comparisons:

1. Ex-PVP vs. all wild accessions
2. Ex-PVP vs. wild1
   Ex-PVP vs. wild2
3. Wild1 vs. wild2

Each comparison was assessed for selective sweep signals using:

- FST (fixation index):
  Calculated using VCFtools in 100 kb windows with 10 kb steps. Windows with FST > 0.3 were considered strongly differentiated, representing likely targets of directional selection.
- π-ratio (nucleotide diversity):
  We computed π for each population independently using VCFtools and calculated the π ExPVP / π Wild ratio. A π-ratio < 0.5 indicated local reductions in diversity among ex-PVPs, suggesting recent or ongoing selection in cultivated lines.
- XP-CLR (Cross-Population Composite Likelihood Ratio):
  XP-CLR (v1.1.2) was applied using 100 kb windows with 10 kb steps and recombination rates estimated from a genetic map. Windows in the top 1% of XP-CLR scores were classified as candidate regions under selection.

To improve specificity, we intersected sweep candidates across methods. Regions overlapping in FST & XP-CLR, or FST & π-ratio & XP-CLR, were retained as high-confidence sweeps, filtering out noise from any single approach.
Genes within ±100 kb of sweep windows were extracted using PyRanges and cross-referenced with the 'BTx623' v5.1 annotation. These gene sets were functionally annotated via eggNOG-mapper and tested for GO term enrichment using the GOATOOLS package. We used a background of all 'BTx623' genes with GO annotations, and adjusted p-values using Benjamini-Hochberg FDR correction.

Enrichment analysis revealed functional themes consistent with domestication and improvement. Candidate sweep regions were enriched for:

- Auxin transport, seed dormancy, and amino acid biosynthesis – pathways implicated in plant architecture, reproductive timing, and seed development.
- Innate immune signaling and xenobiotic detoxification, suggesting shifts in defense strategies during breeding.
- Phospholipase and RNA export activity, indicative of metabolic rewiring under agronomic selection pressures.

In addition to canonical loci, such as SHATTERING1 (SH1), MATURITY1 (MA1), and SORGHUM GRAIN SIZE 3 (SBGS3), we also identified sweeps overlapping circadian and flowering regulators such as PSEUDO-RESPONSE REGULATOR 7 (PRR7), PHOTOTROPIN 1 (PHOT1), CONSTANS (CO), and VERNALIZATION 3B (VRN3B), as well as metabolic integrators like TARGET OF RAPAMYCIN (TOR), REGULATORY-ASSOCIATED PROTEIN OF TOR 1A (RAPTORA), and XAP5 CIRCADIAN TIMEKEEPER (XCT) [12].

Our approach highlights the power of combining population structure inference with multilayered selection metrics to dissect the genomic architecture of domestication and breeding. By filtering for concordant signals across methods and anchoring functional insights in GO enrichment, we prioritized biologically relevant loci for further investigation.

**Circadian and Photoperiodic Gene Networks Underlying Adaptation in Sorghum**

Circadian rhythms in sorghum exhibit regulatory complexity comparable to that of well-studied model plants, integrating light, temperature, and hormonal cues to coordinate key developmental processes [13–16]. In the morning, the *SHAQKYF-type MYB* transcription factor *LATE ELONGATED HYPOCOTYL (LHY)* is activated by *LIGHT-REGULATED WD1 (LWD1)* and *TEOSINTE BRANCHED1/CYCLOIDEA/PCF (TCP)* transcription factors. *LHY* represses the expression of midday and evening genes, including *PSEUDO-RESPONSE REGULATORS (PRR7, PRR9)* and *TIMING OF CAB EXPRESSION 1 (TOC1/PRR1)* [14]. Notably, *LWD* and *TCP* factors also contribute to *PRR* activation, underscoring their dual role in regulating the circadian clock.

By midday, *REVEILLE* transcription factors (*RVE4, RVE8*) and their cofactors *NIGHT LIGHT-INDUCIBLE AND CLOCK-REGULATED* proteins (*LNK1, LNK2*) promote the expression of *PRRs* and the evening complex genes: *EARLY FLOWERING 3 (ELF3), ELF4*, and *LUX ARRHYTHMO (LUX)*. These evening genes are expressed at night and repress morning-expressed genes, forming a feedback loop that stabilizes daily circadian oscillations.

After dusk, the blue-light photoreceptor *ZEITLUPE (ZTL)*—which contains a *LOV (Light, Oxygen, Voltage)* domain—interacts with *GIGANTEA (GI)* to target *PRR5* and *TOC1* for degradation, linking environmental light cues to post-translational regulation of clock components.

In parallel, blue light signaling also influences flowering time by modulating *CONSTANS (CO)* and *FLOWERING LOCUS T (FT)* expression, while *PHYTOCHROME B (PHYB)* perceives red light and regulates growth through *PHYTOCHROME-INTERACTING FACTORS (PIFs)*.

At night, *COLD-REGULATED* genes (*COR27* and *COR28*) help integrate temperature cues by suppressing *ELONGATED HYPOCOTYL 5 (HY5)* activity. Natural variation in core clock genes—particularly *PRR*, *GI*, and *ELF3*—has been associated with adaptation to temperate climates, through changes in photoperiod sensitivity and flowering time [17]. Furthermore, circadian gating of stomatal activity may enhance water-use efficiency and influence herbicide uptake, underscoring the clock's potential applications in precision agriculture and climate-resilient crop design [15].

### Supplementary Figures

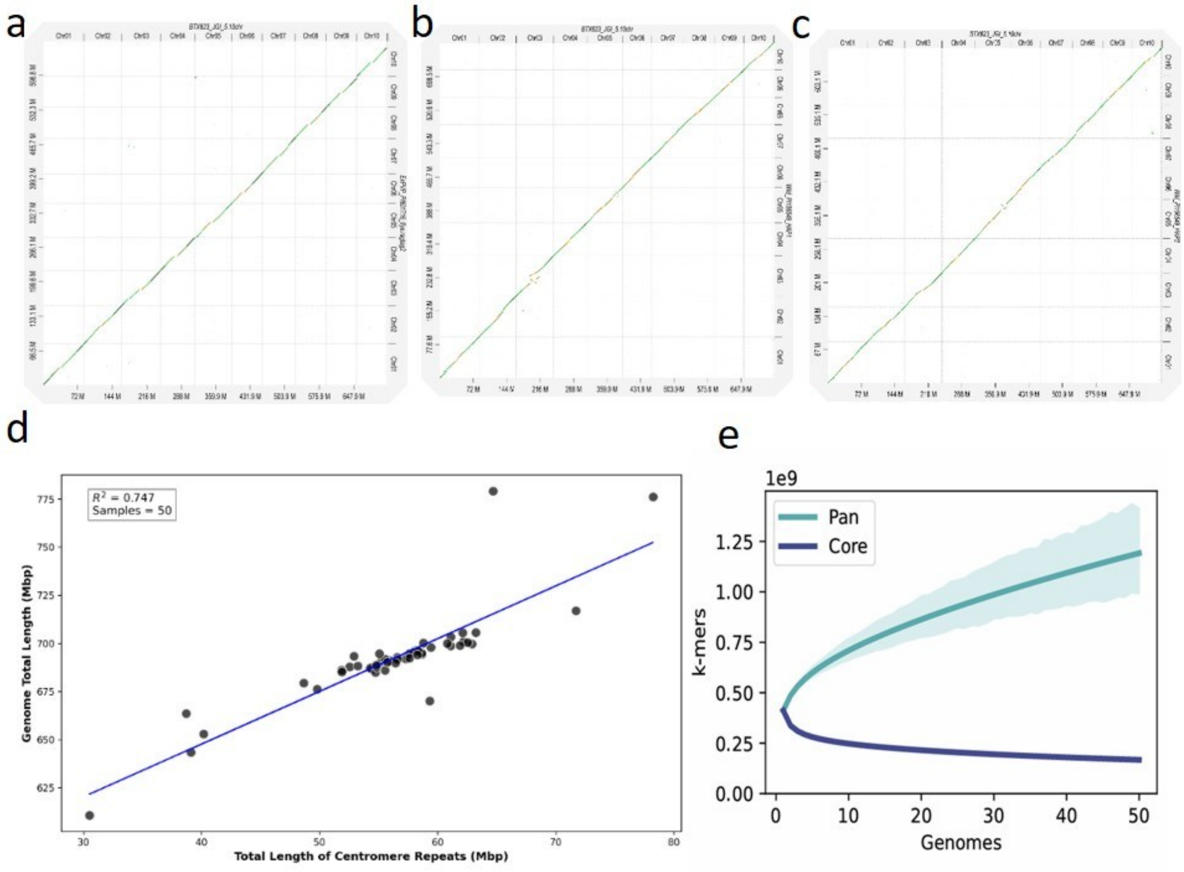


**Supplementary Fig. 1. High-Quality Assemblies Reveal Conserved Synteny and Genome Size Variation Across the Sorghum Pangenome.**

**a–c**, Synteny comparisons between BTx623 (v5) and three assemblies: Ex-PVP PI601716 (a), wild PI156549 haplotype 1 (b), and haplotype 2 (c). Diagonal lines indicate conserved syntenic regions; color intensity reflects sequence identity. **d**, Positive correlation ($R^2$ = 0.747) between centromeric tandem repeat length and genome size across accessions. **e**, PanKmer collector's curve showing continued accumulation of novel k-mers, supporting an open sorghum pangenome. Source data are provided in the Source Data file.

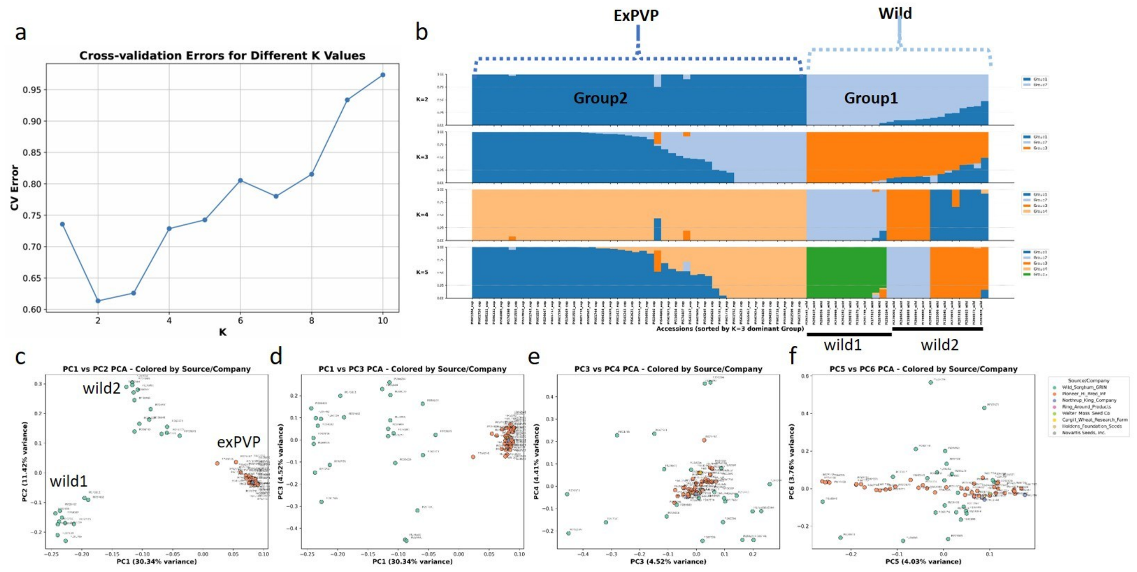


**Supplementary Fig. 2. Genetic Structure and Admixture in Ex-PVP and Wild Sorghum Accessions.**
**a**, Cross-validation errors for different K values, identifying K = 2 as the optimal number of ancestral populations. **b**, Stacked bar plots showing the inferred population structure of 71 sorghum accessions at K = 2, 3, 4, and 5. Colors indicate the proportion of ancestry from each inferred cluster. Accessions are ordered across all K values based on their dominant cluster assignment at K = 3 to facilitate direct comparison. **c**, Principal component analysis (PCA) of 71 sorghum accessions showing PC1 (30.34%) versus PC2 (11.42%). Wild accessions form two major clusters (wild 1 and wild 2), while the ex-PVP accessions form a distinct cluster. **d**, PCA plot showing PC1 (30.34%) versus PC3 (4.52%), further resolving the separation among the three groups identified in the pangenome. **e**, PC3 (4.52%) versus PC4 (4.41%). **f**, PC5 (4.03%) versus PC6 (3.76%). Different dot colors represent the accession source/company, as shown in the legend. Source data are provided in the Source Data file.

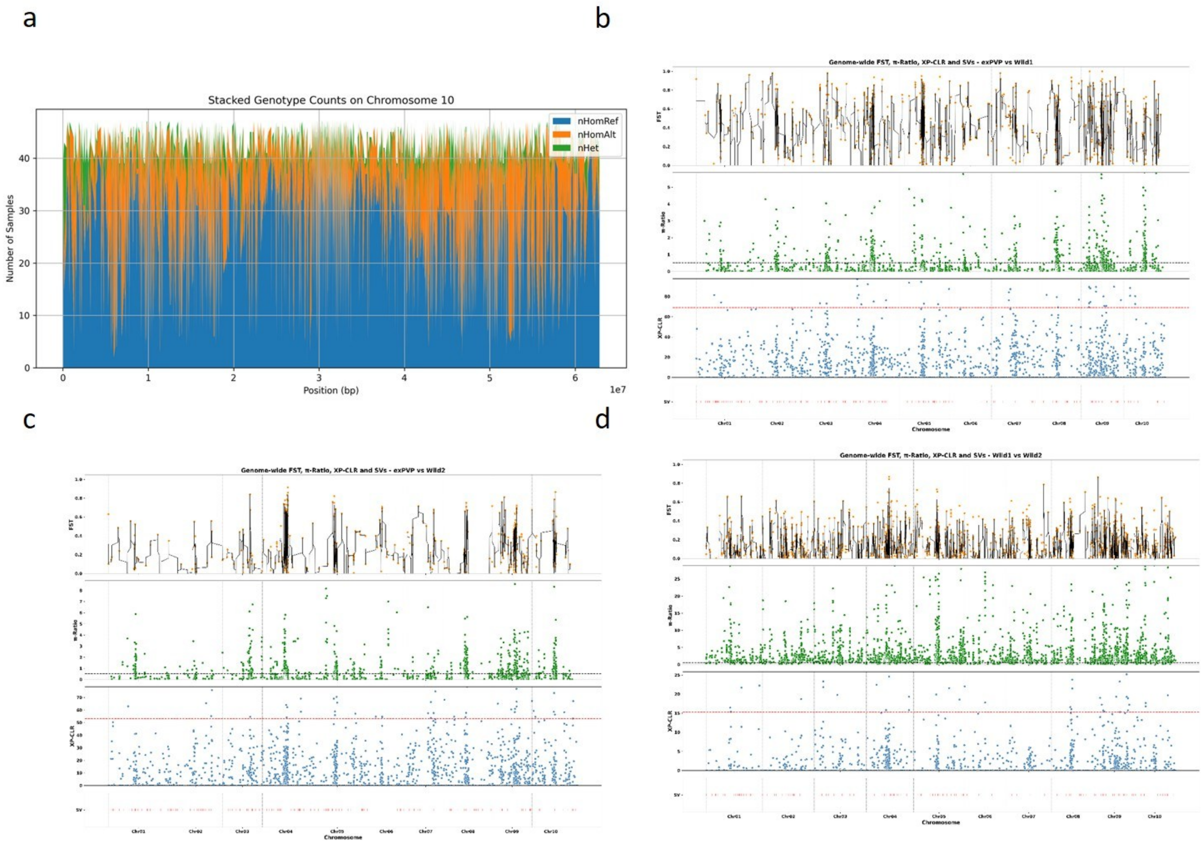


**Supplementary Fig. 3. Zygosity Patterns and Selection Signals Across Sorghum Chromosomes.**

**a**, Zygosity distribution across the 10 sorghum chromosomes. Blue bars represent the number of samples homozygous for the reference allele, orange bars indicate samples homozygous for the alternate allele, and green bars show heterozygous samples. **b**, Genome-wide selective sweep signals (FST, π, XP-CLR) comparing wild group 1 (see Supplementary Fig. 2c) versus 46 ex-PVP accessions. **c**, Selective sweep signals (FST, π, XP-CLR) comparing wild group 2 versus 46 ex-PVP accessions. **d**, Selective sweep signals between wild group 1 and wild group 2. The bottom track in panels **b–d** shows large structural variants identified across the chromosomes. Source data are provided in the Source Data file.

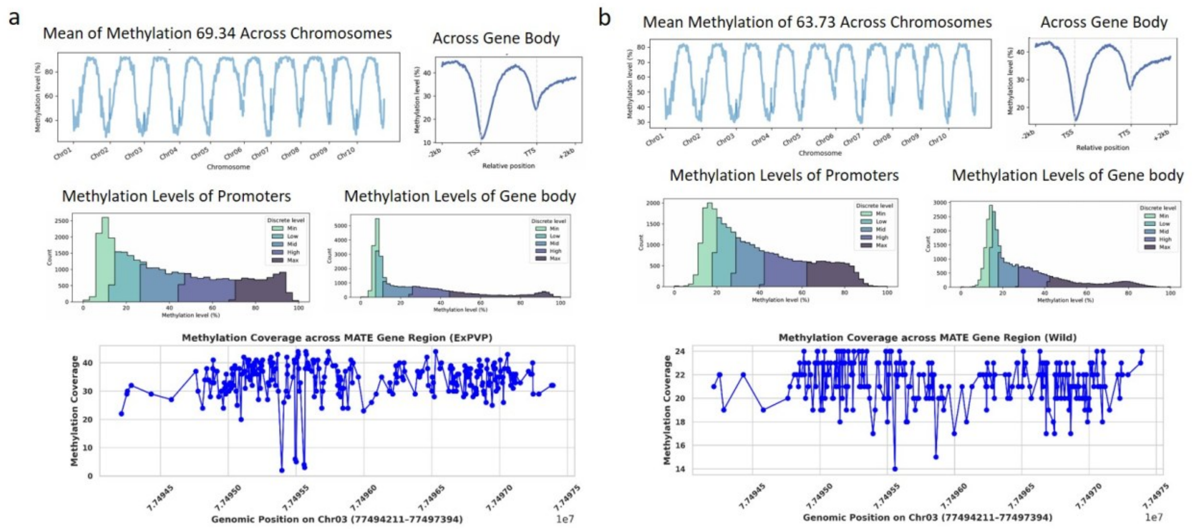


**Supplementary Fig. 4: Genome-Wide and Gene-Body Methylation Profiles of Ex-PVP and Wild Sorghum Accessions.**

**a**, Genome-wide methylation pattern of a representative ex-PVP line (PI562625) with a mean methylation level of 69.34%. **b**, Methylation pattern of a representative wild accession (PI156549) with a mean methylation level of 63.73%. In both panels, line plots illustrate DNA methylation levels across all 10 sorghum chromosomes, promoter and gene body regions, and across the *multidrug and toxic compound extrusion (MATE)* gene on chromosome 3 (Chr03:77,494,151–77,497,397). Notably, segments of the promoter regions show elevated methylation, and differences between cultivated and wild lines are evident across genomic contexts. Source data are provided in the Source Data file.

### Supplementary Tables

**Supplementary Table 1.** Sorghum Pangenome Accessions, Sequencing Platforms, and Assembly Statistics: https://doi.org/10.6084/m9.figshare.29261795.v4

**Supplementary Table 2.** BUSCO Completeness Statistics for Sorghum Genome Assemblies and Predicted Gene Sets

| | **Assembly level** | | | | | | **Protein level** | | | | | |
|---|---|---|---|---|---|---|---|---|---|---|---|---|
| **Accession** | **Complete %** | **Single %** | **Duplicated%** | **Fragmented %** | **Missing %** | **Total BUSCOs** | **Complete %** | **Single %** | **Duplicated%** | **Fragmented %** | **Missing %** | **Total BUSCOs** |
| BTX623 | 99.10% | 96.70% | 2.40% | 0.60% | 0.40% | 1614 | 98.10% | 95.70% | 2.40% | 1.30% | 0.60% | 1614 |
| ExPVP_PI543243 | 99.00% | 96.80% | 2.20% | 0.60% | 0.40% | 1614 | 98.30% | 95.80% | 2.40% | 1.20% | 0.60% | 1614 |
| ExPVP_PI543246 | 99.00% | 96.80% | 2.20% | 0.60% | 0.40% | 1614 | 98.10% | 95.60% | 2.50% | 1.40% | 0.60% | 1614 |
| ExPVP_PI543247 | 98.90% | 96.60% | 2.40% | 0.60% | 0.40% | 1614 | 98.50% | 96.00% | 2.50% | 0.90% | 0.60% | 1614 |
| ExPVP_PI544069 | 98.90% | 96.50% | 2.40% | 0.60% | 0.40% | 1614 | 98.00% | 95.50% | 2.50% | 1.40% | 0.60% | 1614 |
| ExPVP_PI554646 | 98.90% | 96.60% | 2.30% | 0.70% | 0.40% | 1614 | 95.00% | 92.80% | 2.20% | 3.70% | 1.30% | 1614 |
| ExPVP_PI554647 | 98.90% | 96.50% | 2.50% | 0.70% | 0.40% | 1614 | 98.10% | 95.70% | 2.40% | 1.30% | 0.60% | 1614 |
| ExPVP_PI554648 | 98.90% | 96.50% | 2.40% | 0.70% | 0.40% | 1614 | 97.80% | 95.40% | 2.40% | 1.50% | 0.70% | 1614 |
| ExPVP_PI554649 | 99.00% | 96.70% | 2.40% | 0.60% | 0.40% | 1614 | 98.30% | 95.80% | 2.40% | 1.20% | 0.60% | 1614 |
| ExPVP_PI554650 | 98.90% | 96.50% | 2.50% | 0.70% | 0.40% | 1614 | 98.00% | 95.50% | 2.50% | 1.50% | 0.60% | 1614 |
| ExPVP_PI554652 | 98.90% | 96.60% | 2.40% | 0.70% | 0.40% | 1614 | 98.20% | 95.60% | 2.60% | 1.20% | 0.60% | 1614 |
| ExPVP_PI554654 | 98.90% | 96.50% | 2.40% | 0.70% | 0.40% | 1614 | 98.00% | 95.50% | 2.50% | 1.20% | 0.70% | 1614 |

| | Assembly level | | | | | | Protein level | | | | | |
|---|---|---|---|---|---|---|---|---|---|---|---|---|
| **Accession** | **Complete %** | **Single %** | **Duplicated%** | **Fragmented %** | **Missing %** | **Total BUSCOs** | **Complete %** | **Single %** | **Duplicated%** | **Fragmented %** | **Missing %** | **Total BUSCOs** |
| ExPVP_PI555457 | 98.90% | 96.50% | 2.40% | 0.70% | 0.40% | 1614 | 98.00% | 95.40% | 2.70% | 1.30% | 0.70% | 1614 |
| ExPVP_PI561926 | 98.90% | 96.60% | 2.40% | 0.60% | 0.40% | 1614 | 98.20% | 95.90% | 2.30% | 1.20% | 0.60% | 1614 |
| ExPVP_PI562621 | 98.90% | 96.10% | 2.80% | 0.70% | 0.40% | 1614 | 98.00% | 95.00% | 3.00% | 1.40% | 0.60% | 1614 |
| ExPVP_PI562622 | 98.90% | 96.60% | 2.40% | 0.60% | 0.40% | 1614 | 98.10% | 95.50% | 2.60% | 1.20% | 0.70% | 1614 |
| ExPVP_PI562623 | 99.00% | 96.60% | 2.40% | 0.60% | 0.40% | 1614 | 98.00% | 95.50% | 2.40% | 1.20% | 0.80% | 1614 |
| ExPVP_PI562624 | 99.10% | 96.80% | 2.30% | 0.60% | 0.40% | 1614 | 98.30% | 95.80% | 2.50% | 1.20% | 0.40% | 1614 |
| ExPVP_PI562625 | 98.90% | 96.50% | 2.40% | 0.70% | 0.40% | 1614 | 98.00% | 95.50% | 2.50% | 1.30% | 0.70% | 1614 |
| ExPVP_PI564085 | 99.00% | 96.70% | 2.40% | 0.60% | 0.40% | 1614 | 97.90% | 95.40% | 2.50% | 1.50% | 0.60% | 1614 |
| ExPVP_PI574398 | 99.10% | 96.70% | 2.40% | 0.60% | 0.40% | 1614 | 98.30% | 95.80% | 2.50% | 1.20% | 0.60% | 1614 |
| ExPVP_PI574406 | 98.90% | 96.50% | 2.40% | 0.70% | 0.40% | 1614 | 97.60% | 95.10% | 2.50% | 1.60% | 0.80% | 1614 |
| ExPVP_PI574407 | 98.90% | 96.30% | 2.60% | 0.70% | 0.40% | 1614 | 98.20% | 95.40% | 2.80% | 1.20% | 0.60% | 1614 |
| ExPVP_PI594354 | 98.90% | 96.40% | 2.50% | 0.70% | 0.40% | 1614 | 98.30% | 95.70% | 2.50% | 1.10% | 0.60% | 1614 |
| ExPVP_PI594355 | 98.90% | 96.50% | 2.40% | 0.70% | 0.40% | 1614 | 97.70% | 95.20% | 2.50% | 1.70% | 0.60% | 1614 |
| ExPVP_PI595221 | 98.90% | 96.50% | 2.40% | 0.70% | 0.40% | 1614 | 98.30% | 96.20% | 2.20% | 1.10% | 0.60% | 1614 |

| | Assembly level | | | | | | Protein level | | | | | |
|---|---|---|---|---|---|---|---|---|---|---|---|---|
| Accession | Complete % | Single % | Duplicated% | Fragmented % | Missing % | Total BUSCOs | Complete % | Single % | Duplicated% | Fragmented % | Missing % | Total BUSCOs |
| ExPVP_PI596332 | 98.90% | 96.50% | 2.40% | 0.70% | 0.40% | 1614 | 98.40% | 95.80% | 2.60% | 1.20% | 0.40% | 1614 |
| ExPVP_PI596567 | 98.80% | 96.40% | 2.40% | 0.70% | 0.50% | 1614 | 98.00% | 95.60% | 2.40% | 1.40% | 0.70% | 1614 |
| ExPVP_PI601264 | 98.80% | 96.30% | 2.50% | 0.80% | 0.40% | 1614 | 98.40% | 96.00% | 2.40% | 1.20% | 0.40% | 1614 |
| ExPVP_PI601415 | 98.90% | 96.30% | 2.50% | 0.70% | 0.40% | 1614 | 98.40% | 96.00% | 2.40% | 1.10% | 0.60% | 1614 |
| ExPVP_PI601552 | 98.90% | 93.90% | 5.00% | 0.70% | 0.40% | 1614 | 97.00% | 92.40% | 4.60% | 2.10% | 0.90% | 1614 |
| ExPVP_PI601553 | 98.80% | 96.20% | 2.70% | 0.80% | 0.40% | 1614 | 98.20% | 95.50% | 2.70% | 1.10% | 0.70% | 1614 |
| ExPVP_PI601554 | 99.10% | 96.80% | 2.20% | 0.60% | 0.40% | 1614 | 97.80% | 95.50% | 2.30% | 1.40% | 0.80% | 1614 |
| ExPVP_PI601555 | 99.10% | 96.70% | 2.40% | 0.60% | 0.40% | 1614 | 98.00% | 95.60% | 2.40% | 1.40% | 0.60% | 1614 |
| ExPVP_PI601556 | 98.80% | 96.60% | 2.20% | 0.70% | 0.40% | 1614 | 98.00% | 95.60% | 2.40% | 1.40% | 0.60% | 1614 |
| ExPVP_PI601557 | 98.90% | 96.50% | 2.50% | 0.60% | 0.40% | 1614 | 98.30% | 95.80% | 2.50% | 1.20% | 0.60% | 1614 |
| ExPVP_PI601716 | 99.00% | 96.50% | 2.50% | 0.60% | 0.40% | 1614 | 98.10% | 95.50% | 2.50% | 1.40% | 0.60% | 1614 |
| ExPVP_PI601717 | 98.90% | 96.40% | 2.50% | 0.70% | 0.40% | 1614 | 98.00% | 95.50% | 2.50% | 1.40% | 0.70% | 1614 |
| ExPVP_PI601718 | 99.10% | 96.70% | 2.40% | 0.50% | 0.40% | 1614 | 97.10% | 94.60% | 2.50% | 2.10% | 0.70% | 1614 |
| ExPVP_PI601719 | 99.10% | 96.50% | 2.60% | 0.60% | 0.40% | 1614 | 97.80% | 95.20% | 2.50% | 1.70% | 0.60% | 1614 |

| | **Assembly level** | | | | | | **Protein level** | | | | | |
|---|---|---|---|---|---|---|---|---|---|---|---|---|
| **Accession** | **Complete %** | **Single %** | **Duplicated%** | **Fragmented %** | **Missing %** | **Total BUSCOs** | **Complete %** | **Single %** | **Duplicated%** | **Fragmented %** | **Missing %** | **Total BUSCOs** |
| ExPVP_PI601720 | 98.90% | 96.70% | 2.30% | 0.60% | 0.40% | 1614 | 97.90% | 95.60% | 2.30% | 1.50% | 0.60% | 1614 |
| ExPVP_PI601721 | 99.10% | 96.60% | 2.50% | 0.60% | 0.40% | 1614 | 98.00% | 95.40% | 2.60% | 1.50% | 0.50% | 1614 |
| ExPVP_PI601743 | 98.90% | 96.70% | 2.20% | 0.60% | 0.50% | 1614 | 98.20% | 95.80% | 2.40% | 1.20% | 0.60% | 1614 |
| ExPVP_PI601744 | 99.00% | 96.60% | 2.40% | 0.60% | 0.40% | 1614 | 98.00% | 95.50% | 2.50% | 1.50% | 0.60% | 1614 |
| ExPVP_PI601756 | 98.90% | 96.40% | 2.50% | 0.70% | 0.40% | 1614 | 97.90% | 95.40% | 2.50% | 1.50% | 0.60% | 1614 |
| ExPVP_PI602599 | 99.00% | 96.50% | 2.50% | 0.60% | 0.40% | 1614 | 98.50% | 95.90% | 2.50% | 1.00% | 0.60% | 1614 |
| ExPVP_PI602600 | 99.10% | 96.80% | 2.30% | 0.50% | 0.40% | 1614 | 98.00% | 95.50% | 2.50% | 1.40% | 0.60% | 1614 |
| RTx430 | 98.80% | 96.30% | 2.50% | 0.60% | 0.60% | 1614 | 98.20% | 95.70% | 2.50% | 1.20% | 0.60% | 1614 |
| Wild_PI156549_HAP1 | 98.80% | 91.60% | 7.10% | 0.70% | 0.60% | 1614 | 97.70% | 90.50% | 7.20% | 1.40% | 0.90% | 1614 |
| Wild_PI156549_HAP2 | 92.60% | 88.80% | 3.80% | 1.20% | 6.10% | 1614 | 91.70% | 87.60% | 4.10% | 1.70% | 6.60% | 1614 |

**Supplementary Table 3.** PanKmer-Based Adjacency Matrix Showing Pairwise Genomic Distances Among Sorghum Accessions: https://doi.org/10.6084/m9.figshare.29261795.v4
**Supplementary Table 4.** Orthologous Gene Groups Identified in the Sorghum Pangenome: https://doi.org/10.6084/m9.figshare.29261795.v4
**Supplementary Table 5.** KEGG Orthology Annotations for Pangenome-Exclusive Genes in Sorghum: https://doi.org/10.6084/m9.figshare.29261795.v4
**Supplementary Table 6.** Structural Variants Identified in the Sorghum Pangenome Using SVIM-asm and CuteSV:https://doi.org/10.6084/m9.figshare.29261795.v4
**Supplementary Table 7.** Structural Rearrangements Identified by Whole-Genome Alignment Using SyRI:https://doi.org/10.6084/m9.figshare.29261795.v4

**Supplementary Table 8.** Selective sweeps summary

| Comparison | Metric | mRNAs | Genes |
|---|---|---|---|
| exPVP_vs_wild_all | FST | 528 | 392 |
| exPVP_vs_wild_all | PI_ratio | 2736 | 2010 |
| exPVP_vs_wild_all | XPCLR | 3436 | 2519 |
| exPVP_vs_wild_all | FST & XPCLR | 119 | 91 |
| exPVP_vs_wild_all | FST & PI & XPCLR | 71 | 54 |
| exPVP_vs_wild1 | FST | 3112 | 2291 |
| exPVP_vs_wild1 | PI_ratio | 4900 | 3619 |
| exPVP_vs_wild1 | XPCLR | 2788 | 2023 |
| exPVP_vs_wild1 | FST & XPCLR | 531 | 401 |
| exPVP_vs_wild1 | FST & PI & XPCLR | 412 | 298 |
| exPVP_vs_wild2 | FST | 3112 | 2291 |
| exPVP_vs_wild2 | PI_ratio | 4900 | 3619 |
| exPVP_vs_wild2 | XPCLR | 2788 | 2023 |
| exPVP_vs_wild2 | FST & XPCLR | 531 | 401 |
| exPVP_vs_wild2 | FST & PI & XPCLR | 412 | 298 |
| wild1_vs_wild2 | FST | 2366 | 1770 |
| wild1_vs_wild2 | PI_ratio | 1878 | 1389 |

| Comparison | Metric | mRNAs | Genes |
|---|---|---|---|
| wild1_vs_wild2 | XPCLR | 3200 | 2362 |
| wild1_vs_wild2 | FST & XPCLR | 326 | 264 |
| wild1_vs_wild2 | FST & PI & XPCLR | 10 | 9 |

**Supplementary Table 9.** Selective Sweeps Annotation: https://doi.org/10.6084/m9.figshare.29261795.v4

**Supplementary Table 10.** Enriched Circadian Clock and Flowering Time Genes Within Selective Sweep Regions

| Gene | ID | Transcript | Comparison |
|---|---|---|---|
| PRR7 | Sobic.001G411400 | Sobic.001G411400.1.v5.1 | clock_genes_in_FST_exPVP_vs_wild1_selective_sweeps_go_enrichment |
| PRR7 | Sobic.001G411400 | Sobic.001G411400.2.v5.1 | clock_genes_in_FST_exPVP_vs_wild1_selective_sweeps_go_enrichment |
| CO | Sobic.010G115800 | Sobic.010G115800.1.v5.1 | clock_genes_in_FST_exPVP_vs_wild2_selective_sweeps_go_enrichment |
| CO | Sobic.010G115800 | Sobic.010G115800.1.v5.1 | clock_genes_in_PI_exPVP_vs_wild_all_selective_sweeps_go_enrichment |
| RAPTORa | Sobic.005G008800 | Sobic.005G008800.2.v5.1 | clock_genes_in_PI_exPVP_vs_wild_all_selective_sweeps_go_enrichment |
| VRN2b | Sobic.002G164300 | Sobic.002G164300.1.v5.1 | clock_genes_in_PI_exPVP_vs_wild_all_selective_sweeps_go_enrichment |
| TOR | Sobic.009G109200 | Sobic.009G109200.1.v5.1 | clock_genes_in_PI_exPVP_vs_wild_all_selective_sweeps_go_enrichment |
| VRN2b | Sobic.002G164300 | Sobic.002G164300.1.v5.1 | clock_genes_in_PI_exPVP_vs_wild1_selective_sweeps_go_enrichment |
| CO | Sobic.010G115800 | Sobic.010G115800.1.v5.1 | clock_genes_in_PI_exPVP_vs_wild1_selective_sweeps_go_enrichment |
| RAPTORa | Sobic.005G008800 | Sobic.005G008800.2.v5.1 | clock_genes_in_PI_exPVP_vs_wild1_selective_sweeps_go_enrichment |
| TOR | Sobic.009G109200 | Sobic.009G109200.1.v5.1 | clock_genes_in_PI_exPVP_vs_wild1_selective_sweeps_go_enrichment |

| Gene | ID | Transcript | Comparison |
|---|---|---|---|
| VRN3b | Sobic.003G173032 | Sobic.003G173032.2.v5.1 | clock_genes_in_PI_exPVP_vs_wild1_selective_sweeps_go_enrichment |
| RAPTORa | Sobic.005G008800 | Sobic.005G008800.2.v5.1 | clock_genes_in_PI_exPVP_vs_wild2_selective_sweeps_go_enrichment |
| VRN2b | Sobic.002G164300 | Sobic.002G164300.1.v5.1 | clock_genes_in_PI_exPVP_vs_wild2_selective_sweeps_go_enrichment |
| TOR | Sobic.009G109200 | Sobic.009G109200.1.v5.1 | clock_genes_in_PI_exPVP_vs_wild2_selective_sweeps_go_enrichment |
| FRI | Sobic.001G010500 | Sobic.001G010500.1.v5.1 | clock_genes_in_XPCLR_exPVP_vs_wild_all_selective_sweeps_go_enrichment |
| PHOT1 | Sobic.008G001000 | Sobic.008G001000.1.v5.1 | clock_genes_in_XPCLR_exPVP_vs_wild_all_selective_sweeps_go_enrichment |
| PHOT1 | Sobic.008G001000 | Sobic.008G001000.2.v5.1 | clock_genes_in_XPCLR_exPVP_vs_wild_all_selective_sweeps_go_enrichment |
| PHOT1 | Sobic.008G001000 | Sobic.008G001000.3.v5.1 | clock_genes_in_XPCLR_exPVP_vs_wild_all_selective_sweeps_go_enrichment |
| GDH7_ma6 | Sobic.006G004400 | Sobic.006G004400.3.v5.1 | clock_genes_in_XPCLR_exPVP_vs_wild_all_selective_sweeps_go_enrichment |
| FRI | Sobic.001G010500 | Sobic.001G010500.1.v5.1 | clock_genes_in_XPCLR_exPVP_vs_wild1_selective_sweeps_go_enrichment |
| PHOT1 | Sobic.008G001000 | Sobic.008G001000.1.v5.1 | clock_genes_in_XPCLR_exPVP_vs_wild1_selective_sweeps_go_enrichment |
| PHOT1 | Sobic.008G001000 | Sobic.008G001000.2.v5.1 | clock_genes_in_XPCLR_exPVP_vs_wild1_selective_sweeps_go_enrichment |
| PHOT1 | Sobic.008G001000 | Sobic.008G001000.3.v5.1 | clock_genes_in_XPCLR_exPVP_vs_wild1_selective_sweeps_go_enrichment |
| PHOT1 | Sobic.008G001000 | Sobic.008G001000.1.v5.1 | clock_genes_in_XPCLR_exPVP_vs_wild2_selective_sweeps_go_enrichment |
| PHOT1 | Sobic.008G001000 | Sobic.008G001000.2.v5.1 | clock_genes_in_XPCLR_exPVP_vs_wild2_selective_sweeps_go_enrichment |
| PHOT1 | Sobic.008G001000 | Sobic.008G001000.3.v5.1 | clock_genes_in_XPCLR_exPVP_vs_wild2_selective_sweeps_go_enrichment |
| RAPTORa | Sobic.005G008800 | Sobic.005G008800.2.v5.1 | clock_genes_in_XPCLR_exPVP_vs_wild2_selective_sweeps_go_enrichment |
| XAP5 | Sobic.002G277600 | Sobic.002G277600.1.v5.1 | clock_genes_in_XPCLR_exPVP_vs_wild2_selective_sweeps_go_enrichment |

| Gene | ID | Transcript | Comparison |
|---|---|---|---|
| PRR95 | Sobic.002G275100 | Sobic.002G275100.1.v5.1 | clock_genes_in_XPCLR_exPVP_vs_wild2_selective_sweeps_go_enrichment |
| PRR95 | Sobic.002G275100 | Sobic.002G275100.2.v5.1 | clock_genes_in_XPCLR_exPVP_vs_wild2_selective_sweeps_go_enrichment |
| PRR95 | Sobic.002G275100 | Sobic.002G275100.3.v5.1 | clock_genes_in_XPCLR_exPVP_vs_wild2_selective_sweeps_go_enrichment |
| TOR | Sobic.009G109200 | Sobic.009G109200.1.v5.1 | clock_genes_in_XPCLR_exPVP_vs_wild2_selective_sweeps_go_enrichment |

**Supplementary Table 11.** Orthogroups constructed from 46 ex-PVP sorghum lines and two haplotypes from the wild *Sorghum bicolor* accession PI156549. *Arabidopsis thaliana* (Col-0) and *Brassica napus* (Westar) were included as dicot outgroups, and *Zea mays* (B73) as a monocot closely related to sorghum. https://doi.org/10.6084/m9.figshare.29261795.v4

**Supplementary Table 12.** RNA-Seq Sample Details for Eight Tissues Collected from Two Sorghum Accessions (PI329478 and PI510757)

| Reads Yield | File Names at NCBI | Sample |
|---|---|---|
| 5702634152 | SbicPI329478_20220628.ont_pass_cDNA_RNAseq.R000-401.L001-391.fastq.gz | PI329478_Seedling |
| 8165931731 | SbicPI329478_20220628.ont_pass_cDNA_RNAseq.R000-401.L001-392.fastq.gz | PI329478_3 Leaf |
| 6138222035 | SbicPI329478_20220628.ont_pass_cDNA_RNAseq.R000-401.L001-393.fastq.gz | PI329478_5 Leaf |
| 4952355721 | SbicPI329478_20220628.ont_pass_cDNA_RNAseq.R000-401.L001-394.fastq.gz | PI329478_Tiller |
| 5299935238 | SbicPI329478_20220628.ont_pass_cDNA_RNAseq.R000-401.L001-395.fastq.gz | PI329478_Boot |
| 3007973866 | SbicPI329478_20220628.ont_pass_cDNA_RNAseq.R000-401.L001-396.fastq.gz | PI329478_Panicle w / Anthers |
| 3738997776 | SbicPI329478_20220628.ont_pass_cDNA_RNAseq.R000-401.L001-397.fastq.gz | PI329478_Root |
| 6463696818 | SbicPI329478_20220628.ont_pass_cDNA_RNAseq.R000-401.L001-398.fastq.gz | PI329478_dough stage |
| 8403035872 | SbicPI510757_20220628.ont_pass_cDNA_RNAseq.R000-403.L001-399.fastq.gz | PI510757_Seedling |

| Reads Yield | File Names at NCBI | Sample |
|---|---|---|
| 7895491431 | SbicPI510757_20220628.ont_pass_cDNA_RNAseq.R000-403.L001-400.fastq.gz | PI510757_3 Leaf |
| 6534020349 | SbicPI510757_20220628.ont_pass_cDNA_RNAseq.R000-403.L001-401.fastq.gz | PI510757_5 Leaf |
| 6135053379 | SbicPI510757_20220628.ont_pass_cDNA_RNAseq.R000-403.L001-402.fastq.gz | PI510757_Tiller |
| 5261014082 | SbicPI510757_20220628.ont_pass_cDNA_RNAseq.R000-403.L001-403.fastq.gz | PI510757_Boot |
| 2690305273 | SbicPI510757_20220628.ont_pass_cDNA_RNAseq.R000-403.L001-404.fastq.gz | PI510757_Panicle w /Anthers |
| 4054894721 | SbicPI510757_20220628.ont_pass_cDNA_RNAseq.R000-403.L001-405.fastq.gz | PI510757_Root |
| 5145582225 | SbicPI510757_20220628.ont_pass_cDNA_RNAseq.R000-403.L001-406.fastq.gz | PI510757_dough stage |
